# Novel high-content and open-source image analysis tools for profiling mitochondrial morphology in neurological cell models

**DOI:** 10.1101/2024.08.15.607824

**Authors:** Marcus Y. Chin, David A. Joy, Madhuja Samaddar, Anil Rana, Johann Chow, Takashi Miyamoto, Meredith Calvert

**Affiliations:** Denali Therapeutics Inc., South San Francisco, CA 94080 USA

**Author notes:** Authors contributed equally to this work (Marcus Y. Chin, David A. Joy).

**Keywords:** Mitochondria, mitochondrial morphology, high-content imaging, image analysis, high-throughput screening, neurons, open-source, napari plugin

## Abstract

Mitochondria undergo dynamic morphological changes depending on cellular cues, stress, genetic factors, or disease. The structural complexity and disease-relevance of mitochondria have stimulated efforts to generate image analysis tools for describing mitochondrial morphology for therapeutic development. Using high-content analysis, we measured multiple morphological parameters and employed unbiased feature clustering to identify the most robust pair of texture metrics that described mitochondrial state. Here, we introduce a novel image analysis pipeline to enable rapid and accurate profiling of mitochondrial morphology in various cell types and pharmacological perturbations. We applied a high-content adapted implementation of our tool, MitoProfilerHC, to quantify mitochondrial morphology changes in i) a mammalian cell dose response study and ii) compartment-specific drug effects in primary neurons. Next, we expanded the usability of our pipeline by using napari, a Python-powered image analysis tool, to build an open-source version of MitoProfiler and validated its performance and applicability. In conclusion, we introduce MitoProfiler as both a high-content-based and an open-source method to accurately quantify mitochondrial morphology in cells, which we anticipate to greatly facilitate mechanistic discoveries in mitochondrial biology and disease.

## INTRODUCTION

Mitochondria are dynamic organelles responsible for maintaining metabolic homeostasis and generating energy in a eukaryotic cell. They perform critical biochemical processes such as ATP-production, ROS, fatty acid synthesis and calcium regulation (San-Millán, 2023; Zhang et al., 2022). The cell coordinates these functions by regulating the fusion and fission of mitochondria. These molecular mechanisms ultimately determine mitochondrial distribution, size, and morphology, which change in response to various genetic factors, cellular cues, stress and disease (Chan, 2020). Structurally, the mitochondrion consists of a double membrane decorated by proteins. Mitofusin 1, Mitofusin 2 (MFN1, MFN2) and optic atrophy 1 (OPA1) are GTPases that are key regulators of outer and inner mitochondrial membrane fusion (Cipolat et al., 2004; Santel & Fuller, 2001). Dynamin-related protein 1 (DRP1) is one of the main proteins controlling mitochondrial fission (Smirnova et al., 2001). Mutations in these and other fission and fusion proteins cause early onset neurological disorders that can range in severity. For example, *Mfn2* mutations are causal for Charcot-Marie Tooth neuropathy 2A, a disease that preferentially affects axons of peripheral neurons and clinically manifests as muscle weakness (Cartoni & Martinou, 2009). At the cellular level, *Mfn2* deficiency prevents mitochondrial fusion and causes fragmentation of neuronal mitochondria (Chen et al., 2003; Filadi et al., 2018).

Mitochondrial function and ATP generation is particularly important in the brain due to the high energetic needs of neurons. Numerous past studies have identified important molecular links between mitochondria and sporadic forms of neurodegeneration such as Alzheimer’s disease (AD), Parkinson’s disease (PD), and amyotrophic lateral sclerosis (ALS) (Cabral-Costa & Kowaltowski, 2020; Shields et al., 2021; Yang et al., 2021). In neurodegeneration, fragmentation is considered one of the morphological hallmarks of mitochondrial dysfunction (Knott et al., 2008) and precedes neuronal death. The disease-relevance of specific mitochondrial morphologies has fueled the development of quantitative, image-based assays of mitochondrial dynamics, at scales practical for use in therapeutic screening campaigns.

To date, there are many described image analysis programs for mitochondrial morphology; a noncomprehensive list has been presented in Harwig et al. (Harwig et al., 2018). Many studies utilized a range of imaging modalities including epifluorescence, spinning disk confocal, and super-resolution microscopy on semi-automated systems. Both 2D and 3D image analyses have been developed for custom analytical software such as ImageJ/FIJI, MATLAB and Image Pro Plus, capturing conventional mitochondrial measurements including number, area, length, and aspect ratio. Some programs employed supervised machine learning classification models to cluster cells that exhibit defined morphologies (Harwig et al., 2018). For example, one study classified mitochondrial morphologies in mouse photoreceptor cells using an automated wide-field fluorescence microscope, IN Cell Analyzer 2000 Analyzer (Cytiva) (Leonard et al., 2015). In this example, image segmentation of mitochondrial objects was performed using IN Cell Developer Toolbox 1.9.1, while downstream machine-learning classification was performed on the R platform. More recent phenotypic screens have used confocal based high-content systems, including the Opera Phenix system (Revvity, formerly PerkinElmer) to profile the effect of small molecules, environmental toxicants, and neurodegenerative disease mutations on mitochondrial morphology (Charrasse et al., 2023; Little et al., 2018; Varkuti et al., 2020).

High-content screening and analysis (HCS/HCA) has been foundational to therapeutic discovery, allowing researchers to quickly identify novel targets in both candidate-based and phenotypic pipelines (Chin et al., 2021; Way et al., 2023). Here, we introduce MitoProfilerHC, a novel high-content image analysis pipeline that enables the rapid and accurate profiling of mitochondrial morphology. Unlike previously published image-based assays, MitoProfilerHC uses a combination of texture-based measurements to measure mitochondria in individual cells, identified using a custom-built feature selection pipeline. We employ this pipeline to characterize mitochondrial responses to both genetic and chemical perturbations in a variety of cell types including neurons, and we validate the utility of this tool for quantifying mitochondrial morphologies in HCA accurately and at scale.

Commercially available image analysis tools, while powerful and integrated with popular instrumentation, are often proprietary and therefore are inaccessible to most users. To expand usability for all imaging users, we also developed MitoProfiler, an open-source adaptation of our high-content mitochondrial morphology pipeline. MitoProfiler provides an interactive segmentation interface as a napari plugin that allows users to visualize and tune the segmentation and feature extraction components of the pipeline, in addition to a batch mode that enables processing of whole plate datasets. By providing an open-source tool we hope to further expand the accessibility and utility of our mitochondrial analysis platform to the broader scientific community.

## RESULTS

### Generation of MitoProfilerHC workflow to evaluate mitochondrial morphology

To establish a platform capable of reproducibly resolving complex subcellular phenotypes at scale, we first built our mitochondrial image analysis pipeline on a high-content screening system. Because mitochondrial morphology is complex and dynamic and to demonstrate the versatility of the pipeline, we optimized our assays in live cells derived from multiple origins and subjected to multiple perturbations.

Cells were cultured in 96-well assay plates and labeled with Hoechst 33342, CellMask Green Actin, and MitoTracker Deep Red (MTDR). MTDR is a cell-permeant dye that accumulates on active mitochondria in cells and is conventionally used to measure mitochondrial mass, shape and activity. Cells were imaged on a spinning-disk confocal-based Opera Phenix Plus high-content microscope (Revvity) using either a 40X/NA 1.1 or 63X/NA1.15 water immersion objective lens permitting sampling of a larger number of cells at sufficient resolution within a given image due to its high numerical aperture and larger field of view.

Briefly, multi-channel images were first visually rendered in the Harmony software **(Fig. 1A)** and individual cells labeled with Hoechst 33342 were identified using the ‘Find Nuclei’ building block (**Fig. 1B)**. All subsequent analyses were performed on a per-cell basis; therefore, the nuclei identification step should not be bypassed. The ‘Find Cytoplasm’ building block was used to delineate cell borders **(Fig. 1C)**. The 647-channel image of the mitochondria was acquired and pre-processed using the sliding parabola background subtraction method **(Fig. 1D)**. This step produced a sharper image from which the ‘Find Image Region’ building block segmented mitochondrial signal within the cell cytoplasm **(Fig. 1E)**.

**Figure 1:**
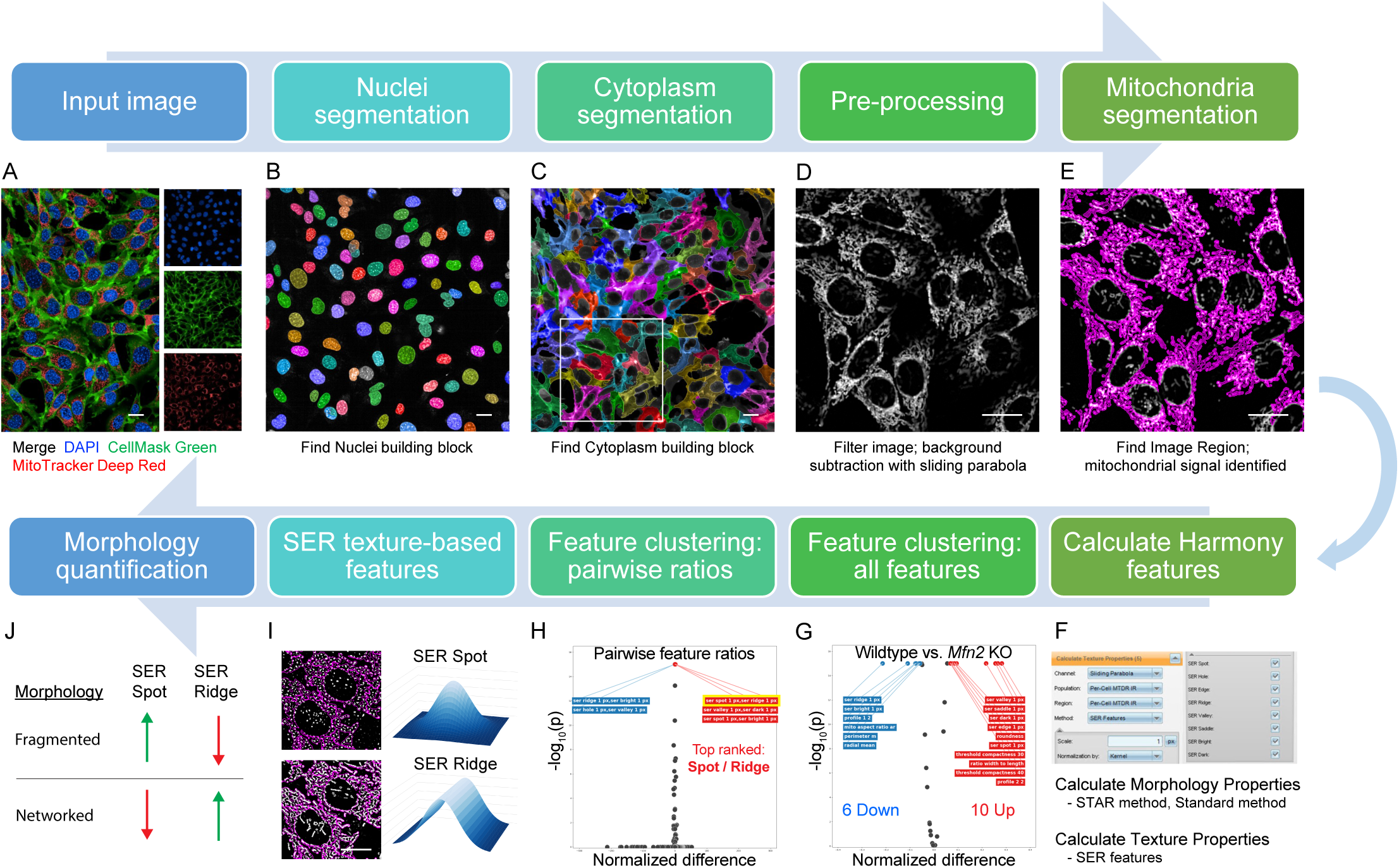
MitoProfilerHC image analysis workflow to evaluate mitochondrial morphology. A) Multi-channel fluorescence input image of WT MEF cells. B) Nuclei segmentation (individual nuclei shown in multi-color). C) Cytoplasm segmentation (individual cells shown in multi-color). Black/white pre-processed image of mitochondria. E) Mitochondrial segmentation (shown in magenta). F) Calculation of Harmony morphology (STAR and Standard) and texture (SER) features. G) Volcano plot of the most discriminative single features that increase with *Mfn2* KO (two-tailed t-test with Holm Sidak correction for multiple comparisons). H) Volcano plot of the most discriminative ratios of features that increase with *Mfn2* KO (two-tailed t-test with Holm Sidak correction for multiple comparisons). I) Representative SER Spot and SER Ridge filtered image (left; individual cells outlined in magenta). Gaussian-derived intensity patterns (right). J) Expected SER Spot and SER Ridge directionality for networked and fragmented mitochondrial morphologies. For all panels, image scale bar = 20 μm.

With mitochondrial structures identified, we next measured morphological and textural features using the ‘Calculate Morphology Properties’ and ‘Calculate Texture Properties’ building blocks, respectively, for each identified mitochondrion **(Fig. 1F)**. The Symmetry-Threshold Compactness-Axial-Radial (STAR) methodology includes a set of properties that classifies phenotypes based on the distribution of intensity within segmented objects. Spot-Edge-Ridge (SER) features are texture filters that are sensitive to different characteristic intensity patterns. Standard morphological properties such as area, roundness, width, and length were also included in the initial protocol development process **(Table 1)**.

**Table 1:** List of Standard Morphology, STAR Morphology, and SER Texture features.

| SER Texture | STAR Morphology |  | Standard Morphology |
| --- | --- | --- | --- |
| SER Spot 1 px | Symmetry 02 | Threshold Compactness 40% | Area (um<sup>2</sup>) |
| SER Hole 1 px | Symmetry 03 | Threshold Compactness 50% | Roundness |
| SER Edge 1 px | Symmetry 04 | Threshold Compactness 60% | Perimeter |
| SER Ridge 1 px | Symmetry 05 | Axial Small Length | Width |
| SER Valley 1 px | Symmetry 12 | Axial Length Ratio | Length |
| SER Saddle 1 px | Symmetry 13 | Radial Mean | Ratio Width to Length |
| SER Bright 1 px | Symmetry 14 | Radial Relative Deviation |  |
| SER Dark 1 px | Symmetry 15 | Profile 1/2 |  |
|  | Threshold Compactness 30% | Profile 2/2 |  |

To generate mitochondrial morphology ground truths in determining which features accurately distinguish mitochondrial states, we acquired images of wildtype (WT) and *Mfn2* deficient mouse embryonic fibroblast (MEF) cells. WT cells under basal conditions were expected to show normal, networked mitochondria while *Mfn2* knockout (KO) cells were expected to show more fragmented morphology. A total set of 33 features were calculated and extracted from mitochondria-segmented images across 9,620 cells (4,810 cells per condition). Feature results were output on a per-cell basis, exported from Harmony, and loaded into a custom-built feature selection pipeline.

Ranking features by largest difference between WT and *Mfn2* KO, we determined that SER features, a family of texture features, were the most robust in distinguishing between fragmented and networked mitochondria **(Table 1).** Ranking individual features by effect size showed that “SER ridge” was the most decreased in *Mfn2* KO compared to WT while “SER valley” was the most increased **(Fig. 1G)**. The most discriminative individual feature was “SER valley” with ∼70.8% classification accuracy when training a logistic regression model on these data (5-fold cross validation with balanced sampling between classes). To further improve this metric, we calculated ratios of all possible pairs of features in the dataset and trained a logistic regression model on the ratio. The ratio of “SER spot” to “SER ridge” was the most discriminative ratio, with ∼75.0% accuracy in cross validation, and was the ratio most increased by *Mfn2* KO **(Fig. 1H) (Table S1)**. SER features detect pixel intensity gradient and curvature; for example, the ‘Spot’ and ‘Ridge’ filters are more sensitive to intensity distributions adopting 3D and 2D Gaussian bell curves, respectively **(Fig. 1I)**. The value of a SER Spot measurement increases in the context of fragmented mitochondria because it is sensitive to round, symmetrical punctate-like structures. Inversely, the value of SER Ridge increases in networked mitochondria exhibiting continuous and elongated structures **(Fig. 1J)**. Together, SER Spot and SER Ridge were identified as the most robust features and were applied in all subsequent morphological analyses performed in cells.

### Validation of MitoProfilerHC in genetic and pharmacological paradigms for mitochondrial disruption

To validate MitoProfilerHC, we quantified mitochondrial morphologies under genetic and pharmacological perturbations in live cells. First, we assessed *Mfn2* KO MEFs, which visually exhibit a striking punctate pattern, indicative of fragmented mitochondria **(Fig. 2A)**. In contrast, unperturbed mitochondria in wildtype (WT) MEFs appear elongated and networked. We first measured the mitochondrial aspect ratio (AR) to provide a benchmark for our analysis in comparison to more traditional methods for quantifying morphology. AR is a proxy for the degree of mitochondrial fusion, defined as the mitochondrial major axis divided by minor axis (i.e. length to width ratio) (Picard et al., 2013). As expected, AR was significantly higher (*p* = 0.0025) in WT cells, reflecting more networked mitochondria **(Fig. 2B)**. Next, we measured mitochondrial morphology in WT and *Mfn2* KO cells using the SER Ridge and Spot textures. Matching the trend in AR, SER Ridge score was significantly higher (*p* < 0.001) in WT cells **(Fig. 2C)**, whereas the SER Spot score was significantly higher (*p* = 0.0023) in *Mfn2* KO cells **(Fig. 2D)**. Overall, these results provided further validation for our high-content analysis method for quantifying both networked and fragmented mitochondria.

**Figure 2:**
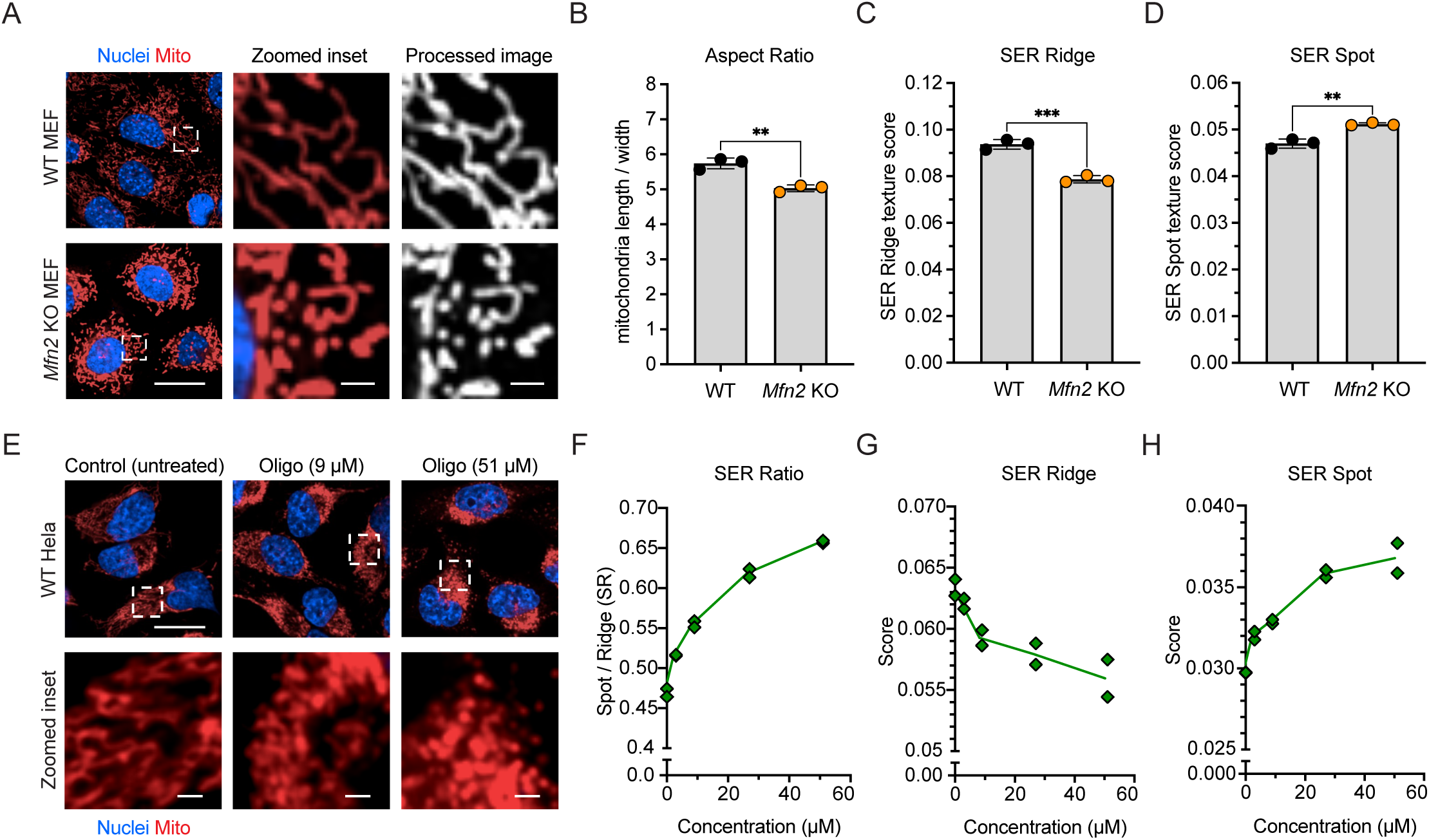
Validating of MitoProfilerHC in genetic and pharmacological paradigms for mitochondrial disruption. A) Representative fluorescence, zoomed inset, and processed images of WT and *Mfn2* KO MEFs. Full-sized image scale bar = 20 μm. Zoomed inset scale bar = 2 μm. B) Mitochondrial aspect ratio (AR) quantification. C) SER Ridge texture quantification. D) SER Spot texture quantification. (B-D) Data points are presented as mean ± SD from three technical replicates; *n* = ∼1,500-2,000 cells per condition group. E) Representative fluorescence images with zoomed inset for WT HeLa cells treated with oligomycin (“Oligo”) from 0 to 51 µM. Full-sized image scale bar = 20 μm. Zoomed inset scale bar = 2 μm. F) SER Ratio (SER Spot/SER Ridge) oligomycin dose-response quantification. G) SER Ridge quantification. H) SER Spot quantification. (F-H) Data points are presented as mean ± SD from two technical replicates; *n* = ∼2,000 cells per condition group. Statistical analysis was performed using two-tailed, unpaired Student’s *t*-test. \*\**p* ≤ 0.01; \*\*\**p* ≤ 0.001; ns = not significant.

The values of SER Spot and SER Ridge consistently yielded an inverse correlation, predictive of either fragmented or networked mitochondrial morphologies, respectively. We further simplified the quantification by taking the ratio of SER Spot-to-SER Ridge, generating a new metric we termed ‘SER Ratio’ (SR). SR thus incorporates both measurements into a single ratiometric value that can be interpreted as the extent of mitochondrial fragmentation. We next sought to demonstrate the applicability of our image analysis pipeline in the context of phenotypic drug discovery using mitochondrial SR. Because mitochondrial dysfunction has a strong link to many human diseases, we moved towards immortalized human cell lines and adapted MitoProfilerHC for Hela cells, a common cell type employed for early-stage drug screening campaigns.

In a comparative study on the effect of pharmacological agents on mitochondrial morphology, oligomycin A was reported to induce robust fragmentation in various cancer cell types (Fu & Lippincott-Schwartz, 2018). Oligomycin is a potent antibiotic that disrupts mitochondrial function by inhibiting proton coupling and ATP synthesis (Hearne et al., 2020; Lardy et al., 1958). In our study, we treated Hela cells for 1.5 hours with oligomycin at a concentration range from 0 to 51 µM, imaged and analyzed mitochondrial morphology using MitoProfilerHC. Mitochondrial fragmentation increased, which is reflected visually **(Fig. 2E)** and through the dose-dependent increase of mitochondrial SR **(Fig. 2F)**. SR values were less variable than its contributing components, SER Ridge and SER Spot **(Figs 2G, H)**, providing a more robust measurement for mitochondrial fragmentation. Importantly, the overall predictive score for mitochondrial SR (∼75%) was higher compared to mitochondrial AR, a commonly used classifier (∼52%) **(Table S1)**. Thus, we were able to demonstrate that MitoProfilerHC provides reduced variability and improved predictive capabilities as compared to commonly used methods and can accurately profile mitochondrial fragmentation in dose-response studies in a human cell line.

### Characterizing drug-induced mitochondrial morphology changes in primary hippocampal neurons using MitoProfilerHC

Neurons are bioenergetically demanding, relying heavily on ATP production and calcium regulation by mitochondria. Mitochondrial dysfunction causes an energetic failure that can acutely induce ischemia (Liu et al., 2018) and has been implicated in several neurological disorders including ALS, Alzheimer’s, and Parkinson’s disease (Cabral-Costa & Kowaltowski, 2020; Norat et al., 2020). To demonstrate its application to drug development in neurological diseases, we adapted the MitoProfilerHC pipeline for quantifying mitochondrial morphology changes under various perturbation conditions in primary hippocampal neurons **(Fig. 3A)**.

**Figure 3:**
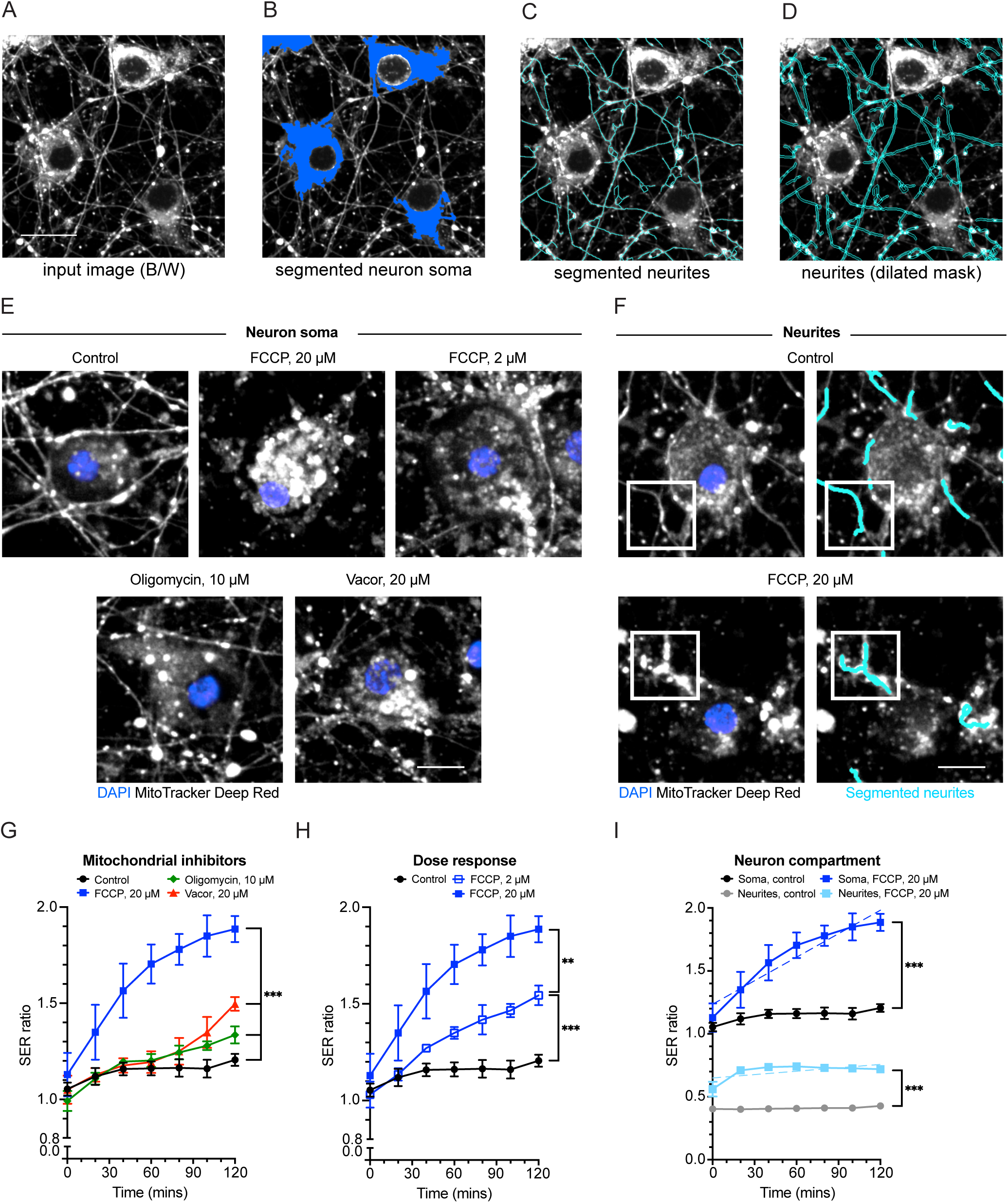
Characterizing drug-induced mitochondrial morphology changes in primary hippocampal neurons using MitoProfilerHC. A) Representative fluorescence input image (pseudo-colored black/white) of mitochondrial channel in primary mouse hippocampal neurons. B) Neuron soma segmentation (blue). C) Neurite segmentation (cyan). D) Dilated neurite mask (cyan). (A-D) All scale bars = 20 μm. E) Representative images of neurons treated with mitochondrial inhibitors, FCCP (2 or 20 µM), oligomycin (10 µM) and Vacor (20 µM) for 2 hrs. F) Representative images of neurons treated with 20 µM FCCP, highlighting mitochondrial fragmentation in neurites (cyan, white boxes). (E-F) All scale bars = 10 μm G) 2 hr time-course quantification comparing 20 µM FCCP (solid blue), 10 µM oligomycin (solid green) and 20 µM Vacor (solid red). H) Dose-response quantification for FCCP, 2 µM (open blue) and 20 µM (solid blue). I) Quantification of SR in neuron soma (solid blue) and neurites (solid light blue) in response to FCCP (20 µM). A simple linear regression comparison was performed on soma and neurites conditions treated with FCCP (dashed lines). (G-I) Data points are presented as mean ± SD from three technical replicates; *n* = ∼500 cells per condition group. Statistical analysis was performed using two-tailed, unpaired Student’s *t*-test. \*\**p* ≤ 0.01; \*\*\**p* ≤ 0.001; ns = not significant.

We further developed the protocol to include segmentation of both neuronal soma and neurite structures **(Fig. 3B, C)**. Given the narrow morphology of neurite processes, we dilated the neurite mask by ∼0.4 µm to ensure detection of the entire structure and its contents **(Fig. 3D)**. Following our previous findings in Hela cells, we treated neurons with various compounds known to disrupt mitochondrial function and induce fragmentation, including oligomycin **(Fig. 3E)**. FCCP, or carbonyl cyanide p-trifluoromethoxyphenylhydrazone, is a commonly used protonophore that uncouples mitochondrial oxidative phosphorylation and inhibits ATP synthesis (Benz & McLaughlin, 1983). It has been extensively characterized in living cells in relation to its inhibitory effect on mitochondrial function. We also used Vacor, a cell permeable precursor of the sterile alpha and TIR motif-containing protein 1 (SARM1) agonist (Loreto et al., 2021), to induce mitochondrial dysfunction. SARM1 plays a crucial role in regulating both axons and dendritic degeneration in hippocampal neurons (Miyamoto et al., 2024; Osterloh et al., 2012) and importantly, is a downstream factor to mitochondrial damage (Summers et al., 2014).

In our comparison between oligomycin, FCCP and Vacor, we found that after two hours of treatment all drugs induced significant mitochondrial fragmentation in neuronal soma, as measured by the SER ratio **(Fig. 3E, G)**. While oligomycin and Vacor fragmented mitochondria slowly and at later timepoints, FCCP induced a much earlier effect that is evident at around 20 minutes post-treatment. Because FCCP had such a rapid and potent effect on mitochondria, we also wanted to test whether our analysis protocol can measure the effect of FCCP in a dose-dependent manner. Indeed, when neurons were treated with FCCP at 2 and 20 µM, we observed a dose-dependent effect **(Fig. 3E, H)**, demonstrating the value of MitoProfilerHC applications requiring *in vitro* dose-response studies of mitochondrial function.

Finally, we investigated whether mitochondria located in different neuronal compartments behave differently in response to pharmacological stress. In neurons that were treated with FCCP at 20 µM, we measured the SR in both soma and neurites **(Fig. 3F, I)**. Compared to control neurites, FCCP induced a significant elevation in SR, suggesting that like soma, mitochondria in neurites also undergo fragmentation in response to chemical stress and indicate that there are compartment-specific responses to perturbation.

### MitoProfiler, an open-source mitochondrial morphology image analysis tool

While MitoProfilerHC was developed on a specific platform using the Harmony software toolkit, we wanted to enable broad usability of our mitochondrial morphology image analysis pipeline to the scientific community. We created MitoProfiler, an open-source version of our previously described methodology using napari, a multi-dimensional image visualization, annotation, and analysis library for Python (Sofroniew et al., 2024). We followed a similar approach as in MitoProfilerHC to extract morphological features from cells. Briefly, we implemented a three-stage segmentation pipeline to first segment cell nuclei followed by cytoplasm and then mitochondria. Next, we implemented a bank of texture features based on the principal curvatures of the mitochondria intensity image similar to the shape index (SI) from (Koenderink and van Doorn, 1992). Finally, we packaged our segmentation and feature extraction code into a napari plugin which can be directly used from an interactive napari session **(Fig. 4A)**.

**Figure 4:**
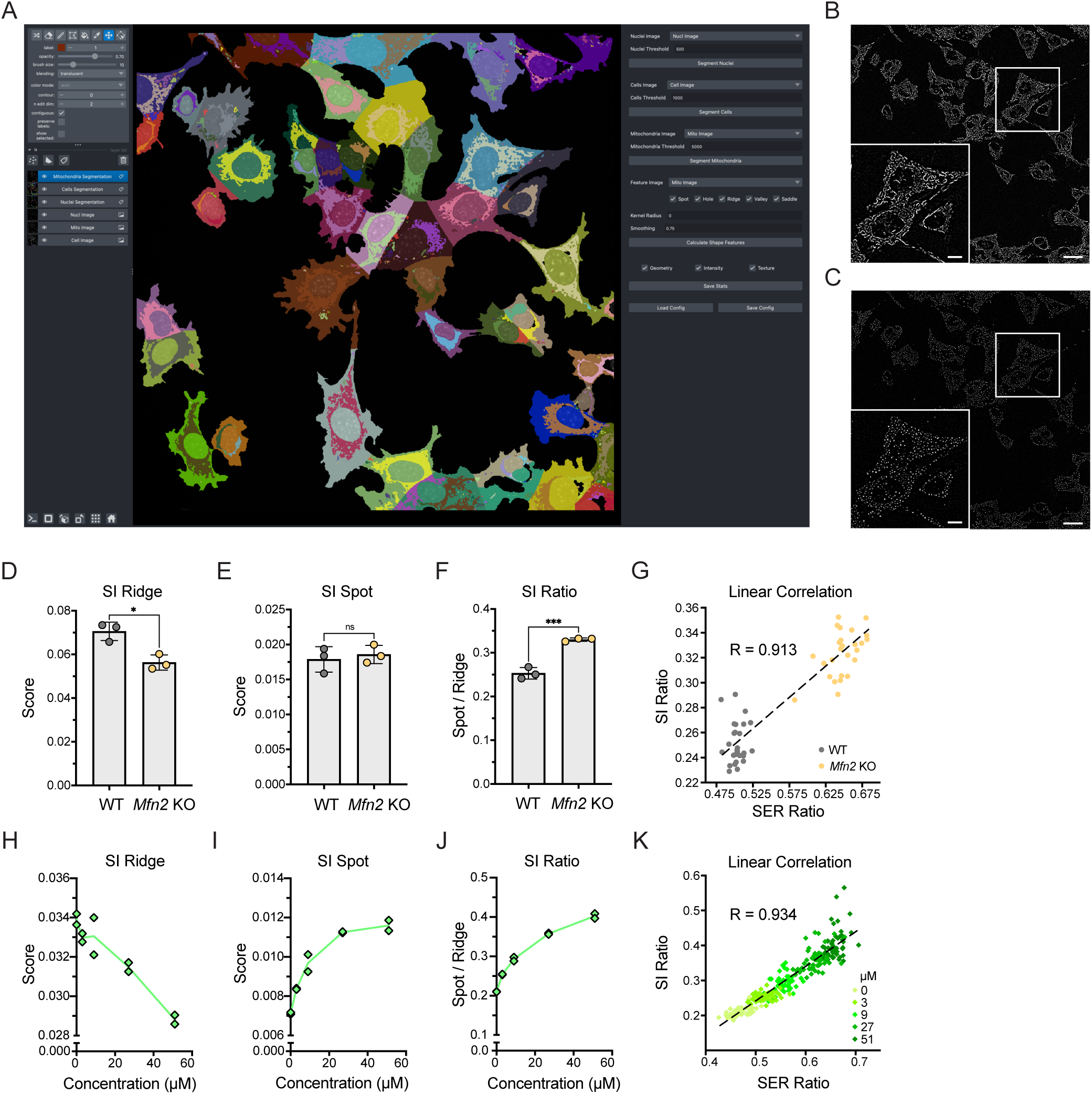
MitoProfiler, an open-source mitochondrial morphology image analysis tool. A) Example session of the MitoProfiler napari plugin analyzing an image of WT MEF cells. Individual cell and mitochondria clusters are colored by cluster identity. Controls to configure individual steps of the MitoProfiler pipeline are shown on the right hand panel. B) SI Ridge and C) SI Spot texture images calculated from the MTDR stained example image with inset to show texture details (scale bar 25 μm for overview image, 10 μm for inset). D) SI Ridge quantification, D) SI Spot quantification and, F) SI Ratio quantification for WT vs *Mfn2* KO MEFs. (D-F) Data points are presented as mean ± SD from three technical replicates; *n* = ∼1,500-2,000 cells per condition group. Statistical analysis was performed using two-tailed, unpaired Student’s *t*-test. *\*p ≤ 0.05*; \*\*\**p* ≤ 0.001; ns = not significant. G) Correlation plot between SI Ratio and SER ratio for *Mfn2* KO experiment with dots individual fields of view colored by WT (gray) or *Mfn2* KO (orange) and a line of best fit (dotted line, slope = 0.501, intercept = 0.000) with Pearson’s correlation coefficient between measurements for each field of view (R = 0.913, p < 1e-10, two-tailed test). H) SI Ridge quantification, I) SI Spot quantification, and J) SI Ratio quantification for WT HeLa cells treated with oligomycin (“Oligo”) from 0 to 51 µM. K) Correlation plot between SI Ratio and SER ratio for treated HeLa cells with dots individual fields of view colored by oligomycin dose and a line of best fit (dotted line, slope = 0.988, intercept = -0.254) with Pearson’s correlation coefficient between measurements for each field of view (R = 0.934, p < 1e-10, two-tailed test).

To demonstrate the utility of our open-source pipeline, we re-segmented the mitochondria in our WT and *Mfn2* deficient (KO) mouse fibroblasts and extracted an SI feature corresponding to ridges (0.75 > SI > 0.25; **Fig. 4B**) and an SI feature corresponding to spots (1.0 > SI > 0.5; **Fig. 4C**) and calculated their per-cell averages (**Fig. 4D, E**, respectively). While the spot feature alone was not significantly different between the two groups (likely due to differences in segmentation between the original Harmony pipeline and our reimplemented pipeline), both the ridge feature alone and the ratio of spot to ridge (SI Ratio) were significantly different between KO and WT **(Fig. 4D-F)**, confirming the robustness of our image texture measures to differences in cell segmentation pipelines. Further, we found good linear correlation between the SER Ratio and our SI Ratio when comparing individual fields of view between the Harmony processed and open-source images (R=0.913; **Fig. 4G**). We next replicated analysis of the dose response study using our shape index-derived spot, ridge, and spot to ridge ratios (**Fig. 4H-J** respectively), validating the mitochondrial feature response from **Fig. 2F-H**. As in the WT vs KO study, the SER Ratio and SI Ratio were well correlated on an individual field of view basis (R=0.934; **Fig. 4K**), indicating that our open-source texture features capture comparable per-image information about mitochondrial fragmentation across experimental modalities. Despite imperfectly reproducing the original Harmony segmentation and featurization pipeline (which is closed source and cannot be directly reimplemented), we believe that our open-source feature extraction pipeline validates the use of spot and ridge texture features as a general measure of microglia fragmentation. The code to calculate these features interactively for a single field of view and as a batch across an imaging study is provided at https://github.com/denalitherapeutics/napari-mito-hcs.

## DISCUSSION

Mitochondrial morphology is highly dynamic and changes rapidly in response to metabolic shifts, cellular perturbation, and various functional impairments in the context of several major diseases. Building upon previous image analysis techniques, we developed a novel high-content image analysis tool, MitoProfilerHC, that robustly quantifies mitochondrial morphology using texture-based measurements in live cells, enabling efficient therapeutic and biomarker screening and development in mitochondrial disorders, cancer, and neurodegeneration. In our study, we provide a comprehensive analysis to determine which texture or morphological feature best predicts relevant mitochondrial phenotypes in healthy and diseased cells with known mitochondrial disruption. MitoProfilerHC is designed for use with the Opera Phenix Plus High-Content System by Revvity, an automated, spinning-disk confocal-based microscope that is widely used in academic and drug discovery research. The MitoProfilerHC analytical pipeline is designed to be easy to use with the Harmony image analysis software that is that is employed for both the acquisition and analysis of images from the Opera Phenix Plus, and our open-source version of MitoProfilerHC can be made compatible with any high content imaging platform that can export images into the standard Tag Image File Format (TIFF). Given the prevalence of high content imaging in both foundational and translational research, we believe this tool will be of broad interest. Additionally, this analytical tool is simple to use and, as our results demonstrate, applicable for use in multiple cell types and assay conditions. Note that although the pipeline is compatible with either of the higher magnification lenses (40X and 63X) that are standard on the Opera Phenix, objectives with lower magnification or resolution (i.e., less than NA=1) are not recommended, due to inadequate resolution of mitochondrial structures.

The SER feature set are texture-based measurements of the spatial distribution of intensity levels in a neighborhood defined through image segmentation strategies and provided as standard metrics within the Harmony image analysis software. We identified two SER texture features – SER Spot and SER Ridge, that when calculated as a ratio, were highly robust in measuring mitochondrial morphology change in drug dose-response studies across multiple cell types, including screenable human cell lines and mature primary neurons. Though previous studies have explored other tools for automating the morphological analysis of mitochondria, to our knowledge there are no published approaches that provide comparable throughput and precision to MitoProfilerHC. One previously published method tracked live mitochondrial movement and classified defined morphologies in response to environmental insult. Image analysis was performed on the CellProfiler software followed by deep learning classification using MATLAB (Zahedi et al., 2018). This method employed traditional confocal imaging and a multi-step image processing and segmentation pipeline and is therefore a substantially lower-throughput approach than that presented here. Another study used the IN Cell Analyzer and IN Cell Developer 1.9.1. image analysis software to segment mitochondria and calculate morphometrics. Subsequently, a machine learning scheme was used in R to classify mitochondrial subtypes based on a priori knowledge of mitochondrial morphology (Leonard et al., 2015). Such morphological classification has been a popular technique to describe complex networks including mitochondria; however, they do not eliminate subjectivity and translatability requires further validation. Additionally, this instrument and associated software is no longer supported by Cytiva Life Sciences (Leonard et al., 2015). Other functionally similar high-content systems include the CellVoyager CV8000 by Yokogawa and ImageXpress Confocal HT.ai by Molecular Devices and it would be interesting to consider adapting our texture-based analytical pipeline for use with their acquisition software. Our tool improves upon previous morphometric-based analyses by enhancing both speed and accuracy of mitochondrial assays performed at-scale under cellular conditions.

Although recent advances in ensemble methods such as deep learning have revolutionized many aspects of image processing in cell biology, classic feature-based approaches such as MitoProfilerHC remain competitive in domains where minimal training data is available, and where computational resources are at a premium such as in high throughput screening campaigns (Chai et al, 2023). Further, while deep learning approaches show impressive performance on classification benchmarks (Natekar et al 2023), establishing methods to explain the “black box” nature of their predictions remains an area of active research (Chai et al 2023, Allen et al 2024, Samek et al 2021). Interpretable feature-based methods such as MitoProfilerHC help build confidence in algorithmic classification of images, both by providing insight into which features of an image are salient to understanding the underlying biology, and by providing confidence that the algorithm can reasonably extrapolate outside of the domain it was initially trained on (Chai et al 2023, Allen et al 2024).

In parallel with our high-content pipeline, we have also developed an open-source version called MitoProfiler that will be made publicly available. Our image analysis work, both in this manuscript and elsewhere has greatly benefitted from the availability of high-quality open-source implementations of image processing algorithms. Building novel analysis algorithms is iterative, and sharing source code across the broader bioimaging community not only enhances efficiency but also data reproducibility as well (Levet et al, 2021). By providing our tool as a plugin to the popular image analysis library napari, we hope to improve the accessibility of our texture analysis method, especially for image analysts who prefer to interact with their data through an open-source graphical user interface (Jamali et al, 2022). Further, we hope to encourage other software developers, especially our colleagues outside of academia, to provide more extensive disclosure of the algorithms that the community relies on to accurately analyze image data.

The image analysis tools that we have developed allowed us to take a deeper dive into investigating mitochondrial function, particularly in the context of neurobiology. Mitochondrial function is crucial in supplying the large bioenergetic demands of neurons (López-Doménech & Kittler, 2023). Its regulation starts during early neuronal development and persists throughout the lifetime of a neuron to ensure survival and protection against neurodegeneration (Rangaraju et al., 2019; Rugarli & Langer, 2012). As such, understanding neuronal response to mitochondrial inhibition has been extensively studied using mitochondrial targeting tool compounds, such as FCCP and oligomycin that both negatively impact the electron transport chain (ETC) and cause subsequent mitochondrial fragmentation. FCCP fragmented mitochondria maximally starting at earlier timepoints and oligomycin induced a milder fragmentation effect at only the highest tested concentration and at later time points. FCCP acts as a rapid protonophore dissipating the proton gradient across mitochondrial membranes, while oligomycin inhibits ATP synthase at the final ETC step. We therefore posit that the degree of ETC disruption is directly correlated with mitochondrial morphology change in neurons.

Similarly, Vacor fragmented mitochondria to a lesser extent compared to FCCP which can potentially be explained by NAD+ depletion by SARM1 agonism and subsequent inhibition of ATP production (Ko et al., 2021; Sato-Yamada et al., 2022). Vacor-mediated activation of SARM1 causes degeneration in all neuronal compartments including cell bodies, axons, and dendrites in primary hippocampal neurons (Miyamoto et al., 2024). Indeed, when we interrogated the effect of mitochondrial inhibition on the neuronal compartment, we observed that mitochondria became fragmented in both neuron soma and neurite processes. We note that the calculated SR is normalized to each region of interest (ROI), thus precluding the direct comparison of absolute SR values between ROIs. In this case, since the overall quantified mitochondrial region was much larger in neurites compared to soma, lower absolute values of SR are seen in soma from both the control and FCCP-treated conditions. Nonetheless, the overall fit of the time course curve between FCCP-treated mitochondria in soma and neurites was also significantly different, suggesting that mitochondria exhibit different dynamics in morphological response to pharmacological perturbation, based on localization. Indeed, mitochondrial localization affects their function and dynamics in the soma, axons, and dendrites. Mitochondria also have been described as having compartment-specific morphologies; for example, mitochondria are densely packed in soma, sparse and rounded in axons, and are larger in dendrites to occupy most of the process (Seager et al., 2020). This result indicates that mitochondria are differentially sensitive to environmental stress depending on the neuronal compartment. Despite compartment differences in function and governing transport mechanisms, MitoProfilerHC was able to quantify differences in morphology between soma and neurites under mitochondrial chemical perturbation.

Overall, we demonstrated the wide utility of the MitoProfilerHC and MitoProfiler tool by interrogating mitochondrial morphology under various *in vitro* cellular assays using a high-content and open-source enabled image analysis. We confirmed the effect of a neurological disease-causing genetic mutation, validated dose-response of various mitochondrial inhibitors, and uncovered compartment-specific changes in mitochondrial morphology that corroborated previous findings. By increasing our understanding of mitochondrial dynamics and morphology using HCS and open-source tools, we hope to greatly facilitate the development of therapeutics targeting mitochondrial diseases.

## MATERIALS AND METHODS

### Cell staining

Prior to live cell imaging, cells were stained with Mitotracker (MitoTracker™ Deep Red FM Dye, Invitrogen, M46753) (MTDR) and Hoechst 33342 (NucBlue™ Live ReadyProbes™, Invitrogen, R37605). The media was removed and a pre-warmed (37°C) staining solution containing MTDR probe (200nM concentration) and Hoechst 33342 probe (75uL per 1ml working concentration) was added to the wells. Cells were incubated at 37°C for 60 minutes. After the staining was complete, the staining solution was replaced with fresh prewarmed media and cells prior to imaging.

### HeLa cells culture and treatment with oligomycin

20,000 HeLa cells were plated per well of a 96 well-plate for a day in 100 µL of DMEM + 10% FBS media. The following day, cells were treated with different concentrations of Oligomycin (Sigma-Aldrich, 75351) for 1.5 hours by adding Oligomycin directly to the cell culture media at the required concentration.

### Mouse embryonic fibroblast cell culture

Mouse embryonic fibroblasts (MEF), WT and *Mitofusin* 2 Knockout (*Mfn2* KO), were cultured in DMEM supplemented with 10% FBS and Penicillin/Streptomycin. 10,000 MEF cells were plated per well of a 96 well-plate for one day. Cells were then live stained with MTDR and Hoechst 33342, as described in the cell staining protocol above.

### Primary mouse hippocampal neurons culture, pharmacological treatment, and staining

Neurons were isolated from mouse embryonic hippocampi (CD-1 strain; Charles River Laboratories) and maintained using NbActiv4 medium (BrainBits; #NB4500) supplemented with Penicillin/Streptomycin (Gibco, #15140122), GlutaMAX (Gibco, #35050061) and 5-fluoro-2-deoxyuridine (Sigma, 50-91-9), as described in (Miyamoto et al., 2024). On day 8 in vitro (DIV 8), neurons were stained with 100 nM MTDR (Invitrogen, #M22426), (1/1000) and NucBlue (Invitrogen, #R37605) for 20 min at 37°C/5% CO2. The mitochondrial inhibitors, Carbonyl cyanide 4-(trifluoromethoxy)phenylhydrazone [FCCP; 2, and 20 µM (MedChemExpress, 370-86-5)], Oligomycin [10 µM; (Sigma, 1404-19-9)], or Vacor [Pyrinuron; 20 µM (ChemService, 53558-25-1)], were added to the stained neurons and imaging was immediately initiated with 20 min time intervals up to 140 min, as described below.

### High-content image acquisition in live cells

All images were acquired on a spinning disk confocal Opera Phenix^™^ Plus High-Content imager (Revvity) using a 4-camera setup (16-bit sCMOS, 6.5 µm pixel size), with two-peak autofocus and 2×2 pixel binning. Environmental controls (37°C/5% CO2) were used for live cell imaging. Fixed wavelength lasers and emission bandpass filters were used to detect fluorophores (Hoechst 33342 E_x_/E_m_: 405/435-480 nm; CellMask E_x_/E_m_: 488/500-550 nm; MTDR E_x_/E_m_: 640/650-760 nm). Acquisition settings were adjusted as needed depending on cell type and density to maximize signal while avoiding saturation and photobleaching. MEFs were imaged on a 40X/1.1 NA water immersion lens (Revvity part number: HH14000422) for Hoechst 33342 (100 ms, 70% power), CellMaskGreen (100 ms, 70% power) and MTDR (100 ms, 80% power) over 9 randomly selected, equally spaced fields with 4 Z-planes (-1.0 to 2.0 µm, 1 µm step size). Hela cells were imaged on a 63X/1.15 NA water immersion lens (Revvity part number: HH14000423) for Hoechst 33342 and MTDR over 27 fields with 4 Z-planes, as detailed. Primary mouse hippocampal neurons were imaged on a 63X/1.15 NA water immersion lens (Revvity part number: HH14000423) with gentler exposure settings to limit phototoxicity (Hoechst 33342: 80 ms, 80%; CellMask Actin: 60 ms, 60%; MTDR: 40 ms, 80%) over 19 fields with 4 Z-planes (-3.0 to 0 µm, 1 µm step size).

### Mitochondrial image analysis, morphology and texture calculation, and data output

The following protocol was built on Harmony 5.2 (Revvity) and applied across multiple cell types. Unless otherwise mentioned, image analysis steps remained consistent across experiments. To achieve even focus, a maximum projection of two planes was taken as the input image. First cells were identified using the ‘Find Nuclei’ building block in the Hoechst 33342 channel (method B, common threshold = 0.0, area > 50 µm^2^, splitting coefficient = 9.8, individual threshold = 0.29, contrast = 0). Next, cytoplasm was identified using ‘Find Cytoplasm’ in the CellMask Green channel (method D, individual threshold = 0.59). The MTDR channel was pre-processed with ‘Filter Image’ (method: sliding parabola, curvature = 50). Mitochondrial signal was segmented using ‘Find Image Region’ (threshold = 0.07, area > 8.75 50 µm^2^) and the mitochondrial morphological and texture metrics were measured. Morphology was calculated using ‘Calculate Morphology’ using two methods: STAR (select Symmetry, Threshold Compactness, Axial, Radial, Profile) and Standard (select Area, Roundness, Perimeter, Length, Width, Width-to-length ratio). ‘Calculate Texture’ was used to measure SER features (method: SER, select SER Spot, Hole, Edge, Ridge, Valley, Saddle, Bright, Dark). Mitochondrial aspect ratio was measured with ‘Calculate Properties’ by taking the formula: mitochondrial per-cell length / per-cell width. Additional steps were included directly after mitochondrial segmentation to identify neurites in primary neuron experiments. The ‘Find Neurites’ building block was used to identify neurite projections attached to neuron cell bodies (Channel: MTDR; Population: Hoechst 33342; Region: Cell; Method: CSIRO Neurite Analysis 2). To quantify the entire neurite, the mask was dilated laterally with ‘Select Region’ (Population: Neurites; Region: Neurite Segment; Method: Resize Region with Outer Border = -2.0 px and Inner Border = INF px). Finally, per-cell results were exported as means per well into Microsoft Excel for further analysis.

### Mitochondrial morphology feature selection

Initial feature selection was performed using scikit-learn v1.3.0 (Pedregosa et al., 2011). First, feature vectors were randomly drawn without replacement for cells in both the WT and KO conditions to create a balanced dataset with 50% WT and 50% KO cells. Next, each feature was mapped to a uniform distribution using the QuantileTransformer in scikit-learn. Features were ranked by effect size (Cohen’s d) and all features with p < 0.01 (two-tailed t-test from scipy v1.11.2 (Virtanen et al., 2020), with the Holm Sidak correction for multiple comparisons from statsmodels v0.14.0 (Seabold et al., 2010) were plotted on a volcano plot. To estimate the accuracy of each feature as a classifier, the data were split into 5 cross validation folds. Each individual feature was used to train a LogisticRegression model with each of the 5 folds (training on 4 out of 5 folds and evaluating on the 5^th^ fold) using scikit-learn with accuracy reported as the average evaluation set score across all 5 folds. All possible combinations of pairs of features were generated and ratios between those pairs were used to train and evaluate a LogisticRegression model as described above.

### Open-source segmentation and feature extraction pipeline

Mitochondria were segmented and assigned to cell masks using a three-phase pipeline. First, the Hoechst-stained nuclei were thresholded and then split into individual nuclei using a watershed transform with a minimum spacing of 3 µm using the watershed segmentation function in scikit-image v0.21.0 (van der Walt et al 2014). Next, CellMaskGreen stained cells were thresholded and then cells containing more than one nucleus were further split by assigning each pixel in a cell mask to the closest nuclei. Finally, MTDR stained mitochondria were thresholded and split into 4-connected components, then further split wherever a mitochondria label crossed a cell boundary using the join segmentations function in scikit-image.

Features were extracted from the mitochondrial intensity image with the following workflow. First, if requested, a parabolic kernel was used to remove background using the rolling_ball function in scikit-image. The kernel was defined as an axisymmetric inverted parabola of height h with the following formula:

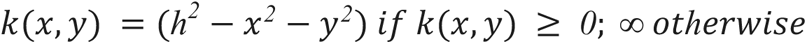

The filtered image was subtracted from the original image and all values less than 0 were set to 0. The image was next smoothed with a gaussian filter to improve the stability of the gradient calculation. Estimates of the first partial derivatives in x and y were calculated using central differences of the smoothed image using the gradient function in numpy v1.24.4 (Harris et al, 2020). The second partial derivatives in xx, xy, and yy were then calculated in a similar manner using the first derivative images. The eigenvalues of the resulting hessian matrix at each point were calculated using the hessian_matrix_eigvals function in scikit-image giving the two principal curvatures at each point with 𝜆*_1_* ≥ 𝜆*_2_*. The shape index from (Koenderink and van Doorn, 1992). was next calculated as:

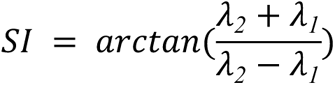

To more closely match the SER images generated by Harmony we selected empirical cutoffs of the shape index approximately twice the width of those given by (Koenderink and van Doorn, 1992). Further, we found that multiplying the shape index by either the first or second principal curvature depending on the texture of interest improved the quality of the resulting image. Specifically, our texture images are defined as:

TI = threshold(SI) * weight

Where:

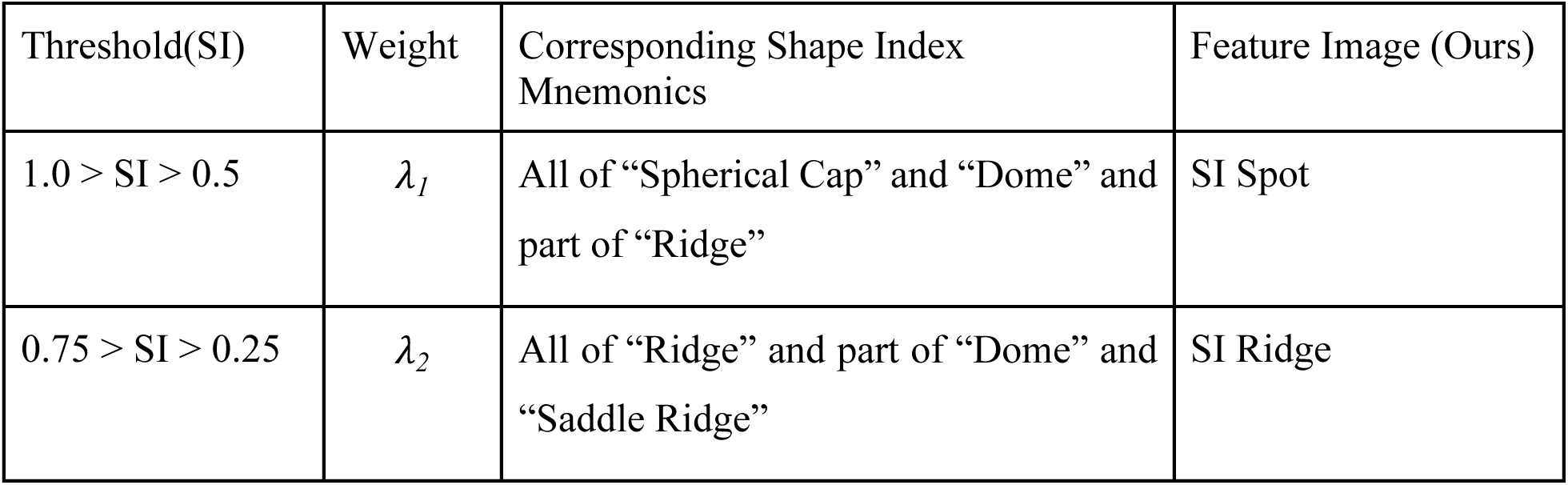
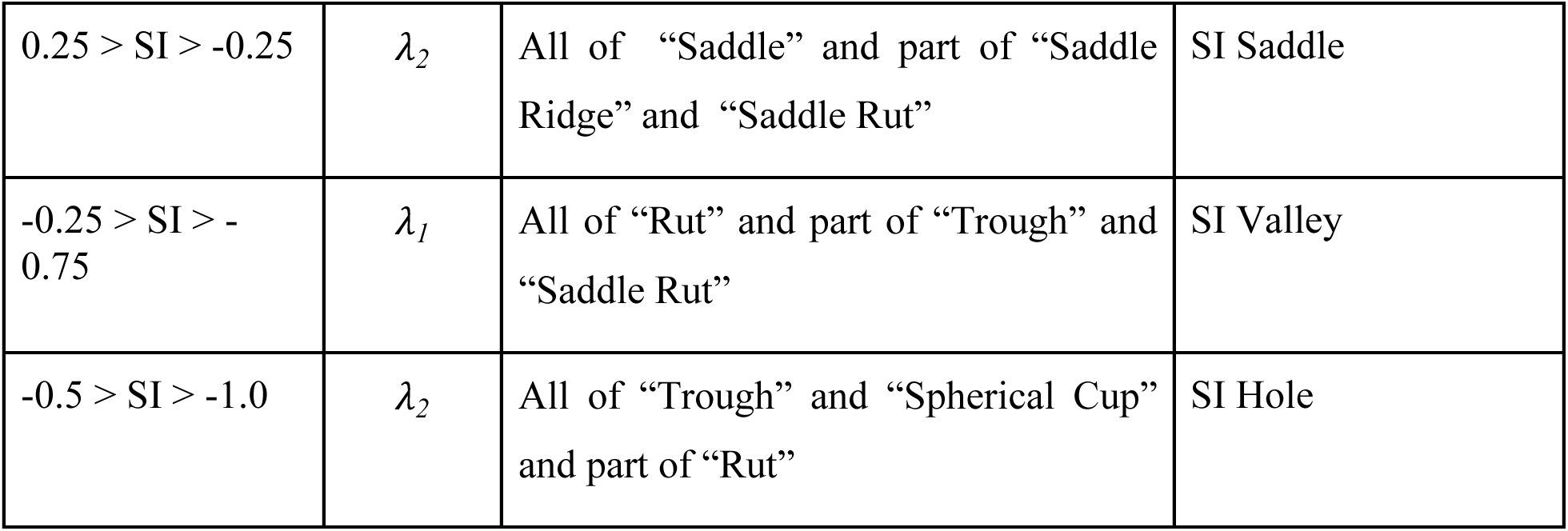

We did not investigate whether this combination of shape index and principal curvature could be extended to approximate the remaining three SER features.

To extract final feature values per-field of view, we used the regionprops_table function in scikit-image to calculate mean values of each feature image within each cell segment. We then calculated averages of the features per-field of view using the group by method in pandas v2.2.0 (McKinney 2010). We calculated the per field of view ratio of SI Spot to SI Ridge, then averaged over all fields of view within a well to get the final values for SI Spot, SI Ridge, and SI Ratio presented in **Fig 4**. Correlation plots were calculated using the per-field of view values for SI Ratio and SER Ratio respectively, fitting a line of best fit using the polyfit function in numpy, then calculating the correlation coefficient and two-sided p-value using the pearsonr function in scipy.

### Data presentation, statistical analysis and illustrations

Data was organized and imported into GraphPad Prism 9 for statistical testing and plotting. A minimum of three wells were imaged per plate and the Number of cells varied depending on the cell type and experiment, as indicated in the Figure Legends. Significant differences between experimental groups were indicated as \**P* < 0.05; \*\**P* < 0.01; \*\*\**P* < 0.001; only *P* < 0.05 was considered as statistically significant. NS, not significant. Schematics were created on Microsoft PowerPoint, SER feature plots were generated on matplotlib v3.8.2 (Hunter et al., 2007), and figures were assembled on Adobe Illustrator.

## ACKNOWLEDGEMENTS

We thank Tanya Weerakkody and Kathryn Monroe for early thoughtful discussions surrounding interpretation of morphological parameters, Lesley Kane for providing insights on mitochondrial biology and developing historical data in cells, Thomas Sandmann for project guidance and review of manuscript, and Robert Graves (Revvity, Inc.) for technical assistance with image analysis inquiries.

## AUTHOR CONTRIBUTIONS

M.Y.C., M.S., M.C. conceived of the study idea and approaches. M.Y.C, M.S., A.R., T.M., M.C. designed experiments. M.Y.C., M.S., A.R. J.C. performed experiments. M.Y.C., M.S., A.R., D.A.J., M.C. analyzed and interpreted data. D.A.J. wrote and tested the code for the open-source pipeline. M.Y.C. and M.C. wrote the manuscript. All authors edited the manuscript.

## DATA AVAILABILITY

All relevant data can be found within the article and its supplementary information.

