## Supplemental Table 1 for "Novel high-content and open-source image analysis tools for profiling mitochondrial morphology in neurological cell models"

**Single Features**

| Features | TrainScore | TestScore |
|---|---|---|
| ser valley 1 px | 0.708627859 | 0.708835759 |
| ser saddle 1 px | 0.704209979 | 0.704054054 |
| ser dark 1 px | 0.683809771 | 0.683471933 |
| ser edge 1 px | 0.665722453 | 0.665904366 |
| ser ridge 1 px | 0.643373181 | 0.643866944 |
| ser spot 1 px | 0.581314969 | 0.581808732 |
| roundness | 0.563799376 | 0.563929314 |
| threshold compactness 30 | 0.563669439 | 0.563201663 |
| threshold compactness 40 | 0.560784823 | 0.560914761 |
| profile 2 2 | 0.560602911 | 0.560706861 |
| threshold compactness 50 | 0.558341996 | 0.558212058 |
| mito aspect ratio ar | 0.556418919 | 0.556237006 |
| ratio width to length | 0.556418919 | 0.556237006 |
| ser bright 1 px | 0.555561331 | 0.555821206 |
| threshold compactness 60 | 0.553638254 | 0.553534304 |
| profile 1 2 | 0.551065489 | 0.551039501 |
| perimeter m | 0.549272349 | 0.5495842 |
| axial small length | 0.543191268 | 0.543347193 |
| length m | 0.541606029 | 0.541164241 |
| radial mean | 0.540150728 | 0.54022869 |
| area m | 0.535265073 | 0.535239085 |
| radial relative deviation | 0.525467775 | 0.525571726 |
| ser hole 1 px | 0.521985447 | 0.521829522 |
| width m | 0.521101871 | 0.521101871 |
| symmetry 03 | 0.519724532 | 0.519438669 |
| symmetry 14 | 0.518892931 | 0.518607069 |
| symmetry 02 | 0.517385655 | 0.517983368 |
| symmetry 05 | 0.515462578 | 0.515592516 |
| symmetry 15 | 0.513253638 | 0.512785863 |
| symmetry 04 | 0.509433472 | 0.509355509 |
| symmetry 13 | 0.508939709 | 0.508939709 |
| symmetry 12 | 0.506964657 | 0.507172557 |
| axial length ratio | 0.501637214 | 0.501559252 |

**Feature Pairs**

| Features | TrainScore | TestScore |
|---|---|---|
| ser spot 1 px,ser ridge 1 px | 0.750623701 | 0.750935551 |
| ser spot 1 px,ser bright 1 px | 0.740072765 | 0.740748441 |
| ser hole 1 px,ser dark 1 px | 0.71764553 | 0.717879418 |
| ser hole 1 px,ser valley 1 px | 0.70210499 | 0.701351351 |
| ser ridge 1 px,ser bright 1 px | 0.695816008 | 0.696673597 |
| ser ridge 1 px,roundness | 0.624012474 | 0.624116424 |
| ser hole 1 px,ser saddle 1 px | 0.610862786 | 0.611018711 |
| profile 2 2,ser ridge 1 px | 0.610446985 | 0.60997921 |
| ser bright 1 px,roundness | 0.592697505 | 0.593347193 |

| | | |
|---|---|---|
| ser edge 1 px,ser saddle 1 px | 0.586798337 | 0.587214137 |
| profile 1 2,roundness | 0.584485447 | 0.584719335 |
| ser valley 1 px,ser saddle 1 px | 0.577182952 | 0.576715177 |
| radial relative deviation,roundness | 0.573284823 | 0.573700624 |
| ser ridge 1 px,ratio width to length | 0.574402287 | 0.573596674 |
| axial small length,ser ridge 1 px | 0.572141372 | 0.571829522 |
| ser edge 1 px,ser valley 1 px | 0.571517672 | 0.570893971 |
| threshold compactness 50,profile 1 2 | 0.566787942 | 0.567255717 |
| threshold compactness 40,profile 1 2 | 0.565982328 | 0.566320166 |
| symmetry 15,roundness | 0.563877339 | 0.565280665 |
| threshold compactness 30,profile 1 2 | 0.564760915 | 0.565176715 |
| axial length ratio,ser ridge 1 px | 0.565462578 | 0.564864865 |
| symmetry 03,roundness | 0.563019751 | 0.562785863 |
| symmetry 13,roundness | 0.5629158 | 0.562681913 |
| symmetry 05,roundness | 0.562733888 | 0.561954262 |
| symmetry 14,roundness | 0.562733888 | 0.561850312 |
| ser valley 1 px,length m | 0.562110187 | 0.561642412 |
| symmetry 13,ser ridge 1 px | 0.56018711 | 0.560602911 |
| ser valley 1 px,mito aspect ratio ar | 0.56049896 | 0.560602911 |
| ser dark 1 px,length m | 0.55997921 | 0.55977131 |
| threshold compactness 60,profile 1 2 | 0.559537422 | 0.559459459 |
| symmetry 04,roundness | 0.558264033 | 0.558939709 |
| symmetry 03,ser ridge 1 px | 0.558419958 | 0.558835759 |
| area m,roundness | 0.558186071 | 0.558731809 |
| ser saddle 1 px,mito aspect ratio ar | 0.55704262 | 0.557380457 |
| ser dark 1 px,mito aspect ratio ar | 0.558523909 | 0.557068607 |
| symmetry 05,ser ridge 1 px | 0.555691268 | 0.556860707 |
| radial mean,roundness | 0.556730769 | 0.556340956 |
| axial small length,roundness | 0.556211019 | 0.555717256 |
| ser valley 1 px,ser dark 1 px | 0.554755717 | 0.554781705 |
| ser edge 1 px,ser dark 1 px | 0.554495842 | 0.554573805 |
| symmetry 14,threshold compactness 30 | 0.554573805 | 0.554365904 |
| symmetry 05,threshold compactness 30 | 0.553482328 | 0.553846154 |
| symmetry 03,threshold compactness 30 | 0.554106029 | 0.553742204 |
| symmetry 12,roundness | 0.554209979 | 0.553638254 |
| profile 2 2,ser bright 1 px | 0.554573805 | 0.553222453 |
| symmetry 15,threshold compactness 30 | 0.552884615 | 0.552806653 |
| symmetry 02,roundness | 0.553040541 | 0.552079002 |
| ser valley 1 px,width m | 0.55231289 | 0.551767152 |
| ser edge 1 px,mito aspect ratio ar | 0.551507277 | 0.551767152 |
| ser hole 1 px,roundness | 0.550883576 | 0.551351351 |
| ser bright 1 px,ratio width to length | 0.550337838 | 0.5504158 |
| ser dark 1 px,width m | 0.549818087 | 0.549272349 |
| ser saddle 1 px,length m | 0.548856549 | 0.548960499 |
| symmetry 13,threshold compactness 30 | 0.548856549 | 0.548856549 |
| symmetry 04,threshold compactness 30 | 0.548882536 | 0.548024948 |
| axial length ratio,roundness | 0.546621622 | 0.547297297 |

| | | |
|---|---|---|
| symmetry 12,threshold compactness 30 | 0.547141372 | 0.546569647 |
| symmetry 14,threshold compactness 40 | 0.545504158 | 0.546049896 |
| symmetry 15,threshold compactness 40 | 0.54495842 | 0.545634096 |
| symmetry 05,threshold compactness 40 | 0.546101871 | 0.545634096 |
| threshold compactness 60,roundness | 0.545218295 | 0.545530146 |
| ser edge 1 px,length m | 0.545816008 | 0.545322245 |
| ser spot 1 px,mito aspect ratio ar | 0.543035343 | 0.542411642 |
| symmetry 04,threshold compactness 40 | 0.541761954 | 0.541891892 |
| symmetry 03,threshold compactness 40 | 0.541865904 | 0.541580042 |
| symmetry 12,threshold compactness 40 | 0.540748441 | 0.541372141 |
| symmetry 02,mito aspect ratio ar | 0.540904366 | 0.540956341 |
| threshold compactness 50,roundness | 0.540722453 | 0.54012474 |
| symmetry 02,threshold compactness 30 | 0.54002079 | 0.53981289 |
| symmetry 02,profile 1 2 | 0.539994802 | 0.53960499 |
| symmetry 13,threshold compactness 40 | 0.539656965 | 0.53950104 |
| symmetry 12,mito aspect ratio ar | 0.538331601 | 0.539189189 |
| threshold compactness 30,radial relative deviation | 0.539007277 | 0.538981289 |
| symmetry 14,threshold compactness 50 | 0.539033264 | 0.538149688 |
| profile 2 2,mito aspect ratio ar | 0.537967775 | 0.537525988 |
| symmetry 12,profile 1 2 | 0.536512474 | 0.537006237 |
| ser hole 1 px,ser edge 1 px | 0.5375 | 0.536070686 |
| ser spot 1 px,roundness | 0.535810811 | 0.535966736 |
| symmetry 05,threshold compactness 50 | 0.536356549 | 0.535966736 |
| profile 1 2,ratio width to length | 0.535706861 | 0.535862786 |
| threshold compactness 40,radial relative deviation | 0.535524948 | 0.534927235 |
| symmetry 02,threshold compactness 40 | 0.536044699 | 0.534927235 |
| ser saddle 1 px,ser dark 1 px | 0.53466736 | 0.534407484 |
| symmetry 03,ratio width to length | 0.533991684 | 0.534303534 |
| axial length ratio,profile 1 2 | 0.533471933 | 0.534095634 |
| profile 2 2,roundness | 0.534225572 | 0.533887734 |
| ser saddle 1 px,width m | 0.533186071 | 0.533575884 |
| symmetry 03,threshold compactness 50 | 0.53277027 | 0.533471933 |
| symmetry 15,threshold compactness 50 | 0.532536383 | 0.532952183 |
| radial relative deviation,ratio width to length | 0.53287422 | 0.532640333 |
| symmetry 04,profile 1 2 | 0.532536383 | 0.532224532 |
| symmetry 04,threshold compactness 50 | 0.53297817 | 0.531808732 |
| symmetry 05,ratio width to length | 0.531185031 | 0.531392931 |
| symmetry 15,mito aspect ratio ar | 0.531418919 | 0.531288981 |
| ser edge 1 px,ser ridge 1 px | 0.532900208 | 0.531185031 |
| roundness,mito aspect ratio ar | 0.531600832 | 0.531081081 |
| threshold compactness 50,mito aspect ratio ar | 0.531860707 | 0.531081081 |
| symmetry 04,mito aspect ratio ar | 0.53266632 | 0.531081081 |
| symmetry 14,mito aspect ratio ar | 0.531366944 | 0.530977131 |
| ser saddle 1 px,ser bright 1 px | 0.53245842 | 0.530665281 |
| threshold compactness 40,mito aspect ratio ar | 0.530977131 | 0.53045738 |
| symmetry 13,threshold compactness 50 | 0.52983368 | 0.53045738 |
| threshold compactness 60,mito aspect ratio ar | 0.531185031 | 0.53024948 |

| | | |
|---|---|---|
| threshold compactness 30,mito aspect ratio ar | 0.531548857 | 0.53024948 |
| ratio width to length,mito aspect ratio ar | 0.531211019 | 0.52972973 |
| threshold compactness 40,roundness | 0.52972973 | 0.52972973 |
| threshold compactness 50,radial relative deviation | 0.528638254 | 0.52962578 |
| symmetry 05,mito aspect ratio ar | 0.531133056 | 0.529209979 |
| symmetry 05,profile 1 2 | 0.52793659 | 0.528482328 |
| threshold compactness 50,threshold compactness 6( | 0.528196466 | 0.528274428 |
| perimeter m,ratio width to length | 0.529054054 | 0.528170478 |
| symmetry 14,ratio width to length | 0.529028067 | 0.528066528 |
| symmetry 15,ratio width to length | 0.528768191 | 0.527962578 |
| symmetry 12,threshold compactness 50 | 0.527468815 | 0.527858628 |
| area m,ratio width to length | 0.528222453 | 0.527650728 |
| symmetry 04,ratio width to length | 0.527053015 | 0.527234927 |
| symmetry 15,profile 1 2 | 0.526923077 | 0.527027027 |
| area m,length m | 0.526663202 | 0.526819127 |
| radial mean,ratio width to length | 0.527286902 | 0.526819127 |
| symmetry 13,profile 1 2 | 0.527156965 | 0.526819127 |
| length m,mito aspect ratio ar | 0.527364865 | 0.526715177 |
| symmetry 13,mito aspect ratio ar | 0.52741684 | 0.526507277 |
| length m,ratio width to length | 0.527884615 | 0.526403326 |
| symmetry 03,profile 1 2 | 0.526715177 | 0.526091476 |
| symmetry 13,ratio width to length | 0.525675676 | 0.525987526 |
| axial small length,ratio width to length | 0.526767152 | 0.525883576 |
| radial relative deviation,mito aspect ratio ar | 0.525727651 | 0.525883576 |
| symmetry 03,mito aspect ratio ar | 0.526715177 | 0.525883576 |
| symmetry 14,profile 1 2 | 0.525805613 | 0.525779626 |
| ser hole 1 px,mito aspect ratio ar | 0.526221414 | 0.525675676 |
| axial length ratio,length m | 0.52531185 | 0.525571726 |
| ser hole 1 px,length m | 0.525883576 | 0.525259875 |
| width m,ratio width to length | 0.5254158 | 0.525051975 |
| radial mean,perimeter m | 0.525363825 | 0.524948025 |
| symmetry 13,length m | 0.524740125 | 0.524636175 |
| width m,mito aspect ratio ar | 0.52468815 | 0.523908524 |
| symmetry 02,symmetry 12 | 0.521881497 | 0.523700624 |
| axial length ratio,mito aspect ratio ar | 0.523778586 | 0.523700624 |
| symmetry 02,threshold compactness 50 | 0.524506237 | 0.523700624 |
| threshold compactness 30,roundness | 0.523466736 | 0.523700624 |
| symmetry 02,ratio width to length | 0.523856549 | 0.523492723 |
| axial length ratio,ser bright 1 px | 0.522141372 | 0.523388773 |
| axial small length,ser bright 1 px | 0.523414761 | 0.523388773 |
| width m,length m | 0.524922037 | 0.523284823 |
| area m,mito aspect ratio ar | 0.523726611 | 0.523284823 |
| axial length ratio,ratio width to length | 0.52279106 | 0.522869023 |
| threshold compactness 40,threshold compactness 6( | 0.522661123 | 0.522141372 |
| axial small length,mito aspect ratio ar | 0.521231809 | 0.521517672 |
| ser hole 1 px,ratio width to length | 0.520919958 | 0.521517672 |
| symmetry 12,ratio width to length | 0.521569647 | 0.521413721 |

| | | |
|---|---|---|
| threshold compactness 60,length m | 0.523440748 | 0.521413721 |
| threshold compactness 50,length m | 0.521647609 | 0.521101871 |
| threshold compactness 40,threshold compactness 5( | 0.521439709 | 0.521101871 |
| threshold compactness 60,radial relative deviation | 0.52027027 | 0.520582121 |
| ser edge 1 px,width m | 0.520426195 | 0.52037422 |
| axial small length,profile 1 2 | 0.519984407 | 0.52006237 |
| symmetry 03,ser bright 1 px | 0.518191268 | 0.51954262 |
| ser dark 1 px,roundness | 0.519594595 | 0.51954262 |
| symmetry 04,radial relative deviation | 0.518581081 | 0.518918919 |
| ser bright 1 px,mito aspect ratio ar | 0.518892931 | 0.518607069 |
| radial mean,profile 1 2 | 0.518373181 | 0.518399168 |
| radial relative deviation,profile 1 2 | 0.518321206 | 0.518191268 |
| roundness,length m | 0.520608108 | 0.517983368 |
| radial mean,mito aspect ratio ar | 0.517177755 | 0.517983368 |
| threshold compactness 30,threshold compactness 6( | 0.517593555 | 0.517671518 |
| symmetry 03,radial relative deviation | 0.516398129 | 0.517255717 |
| symmetry 12,radial relative deviation | 0.51512474 | 0.516943867 |
| axial length ratio,radial relative deviation | 0.516502079 | 0.516735967 |
| symmetry 14,radial mean | 0.516164241 | 0.516632017 |
| symmetry 13,radial relative deviation | 0.515956341 | 0.516112266 |
| threshold compactness 30,threshold compactness 5( | 0.515540541 | 0.516112266 |
| perimeter m,mito aspect ratio ar | 0.516372141 | 0.516008316 |
| ser spot 1 px,length m | 0.519282744 | 0.515384615 |
| profile 1 2,mito aspect ratio ar | 0.516554054 | 0.515176715 |
| symmetry 04,ser ridge 1 px | 0.515072765 | 0.515072765 |
| ser edge 1 px,ser bright 1 px | 0.515904366 | 0.514760915 |
| symmetry 14,radial relative deviation | 0.514267152 | 0.514656965 |
| ser valley 1 px,roundness | 0.514475052 | 0.514656965 |
| threshold compactness 60,ratio width to length | 0.515722453 | 0.514553015 |
| symmetry 02,ser ridge 1 px | 0.516268191 | 0.514345114 |
| threshold compactness 30,ser hole 1 px | 0.514319127 | 0.514241164 |
| profile 2 2,length m | 0.519516632 | 0.514137214 |
| ser spot 1 px,ratio width to length | 0.514319127 | 0.514033264 |
| threshold compactness 30,length m | 0.519334719 | 0.513929314 |
| symmetry 05,radial relative deviation | 0.513877339 | 0.513929314 |
| threshold compactness 60,ser ridge 1 px | 0.516580042 | 0.513929314 |
| radial mean,ser ridge 1 px | 0.51512474 | 0.513825364 |
| threshold compactness 60,ser hole 1 px | 0.514241164 | 0.513825364 |
| threshold compactness 40,length m | 0.519230769 | 0.513825364 |
| symmetry 12,ser ridge 1 px | 0.515722453 | 0.513825364 |
| threshold compactness 40,ser hole 1 px | 0.513981289 | 0.513721414 |
| ser hole 1 px,ser ridge 1 px | 0.517385655 | 0.513617464 |
| symmetry 14,ser ridge 1 px | 0.513539501 | 0.513409563 |
| radial mean,ser valley 1 px | 0.513175676 | 0.513201663 |
| symmetry 15,ser ridge 1 px | 0.514137214 | 0.512785863 |
| ser valley 1 px,perimeter m | 0.513877339 | 0.512474012 |
| threshold compactness 50,ser hole 1 px | 0.511330561 | 0.512266112 |

| | | |
|---|---|---|
| ser dark 1 px,perimeter m | 0.513851351 | 0.512162162 |
| symmetry 15,radial relative deviation | 0.510420998 | 0.511850312 |
| axial small length,width m | 0.511954262 | 0.511850312 |
| threshold compactness 50,ser ridge 1 px | 0.516060291 | 0.511850312 |
| profile 2 2,ratio width to length | 0.511642412 | 0.511538462 |
| axial small length,ser spot 1 px | 0.510836798 | 0.511434511 |
| ser valley 1 px,ser bright 1 px | 0.513747401 | 0.511330561 |
| axial small length,ser edge 1 px | 0.515566528 | 0.511330561 |
| ser hole 1 px,width m | 0.510966736 | 0.511122661 |
| threshold compactness 40,ser ridge 1 px | 0.515254678 | 0.510914761 |
| profile 2 2,ser hole 1 px | 0.511278586 | 0.510602911 |
| threshold compactness 30,ser ridge 1 px | 0.514163202 | 0.51039501 |
| radial mean,ser spot 1 px | 0.510576923 | 0.51008316 |
| symmetry 15,axial length ratio | 0.510005198 | 0.51008316 |
| symmetry 13,axial length ratio | 0.509693347 | 0.50997921 |
| threshold compactness 30,threshold compactness 4( | 0.510862786 | 0.50977131 |
| ser edge 1 px,roundness | 0.509173597 | 0.50966736 |
| radial relative deviation,ser ridge 1 px | 0.513019751 | 0.50956341 |
| ser saddle 1 px,perimeter m | 0.511668399 | 0.509355509 |
| axial length ratio,ser spot 1 px | 0.506626819 | 0.509147609 |
| radial mean,ser dark 1 px | 0.507198545 | 0.508419958 |
| symmetry 02,symmetry 03 | 0.508160083 | 0.508316008 |
| radial relative deviation,ser hole 1 px | 0.507068607 | 0.508316008 |
| symmetry 03,ser spot 1 px | 0.507432432 | 0.508212058 |
| radial mean,ser edge 1 px | 0.511772349 | 0.508212058 |
| axial small length,length m | 0.505873181 | 0.508212058 |
| axial length ratio,ser hole 1 px | 0.50766632 | 0.507900208 |
| axial small length,radial relative deviation | 0.507406445 | 0.507796258 |
| threshold compactness 50,ratio width to length | 0.508393971 | 0.507692308 |
| symmetry 05,ser valley 1 px | 0.506964657 | 0.507484407 |
| symmetry 02,radial relative deviation | 0.506626819 | 0.507484407 |
| symmetry 13,ser spot 1 px | 0.507094595 | 0.507484407 |
| radial relative deviation,ser spot 1 px | 0.507276507 | 0.507380457 |
| symmetry 05,ser spot 1 px | 0.50735447 | 0.507380457 |
| radial mean,radial relative deviation | 0.507198545 | 0.507276507 |
| symmetry 15,ser hole 1 px | 0.503872141 | 0.507172557 |
| symmetry 05,length m | 0.508705821 | 0.507068607 |
| roundness,ratio width to length | 0.506314969 | 0.507068607 |
| profile 1 2,ser ridge 1 px | 0.510680873 | 0.506860707 |
| symmetry 14,ser spot 1 px | 0.506808732 | 0.506652807 |
| ser edge 1 px,perimeter m | 0.509589397 | 0.506444906 |
| symmetry 03,length m | 0.509147609 | 0.506340956 |
| symmetry 12,symmetry 14 | 0.503014553 | 0.506237006 |
| symmetry 15,ser spot 1 px | 0.506860707 | 0.506133056 |
| symmetry 04,ser spot 1 px | 0.507016632 | 0.505821206 |
| symmetry 03,ser hole 1 px | 0.506782744 | 0.505717256 |
| symmetry 02,symmetry 05 | 0.506211019 | 0.505717256 |

| | | |
|---|---|---|
| symmetry 12,ser spot 1 px | 0.505977131 | 0.505405405 |
| symmetry 02,ser spot 1 px | 0.50535343 | 0.505301455 |
| symmetry 12,radial mean | 0.506990644 | 0.505197505 |
| symmetry 05,perimeter m | 0.506237006 | 0.504885655 |
| symmetry 02,radial mean | 0.507588358 | 0.504885655 |
| symmetry 05,ser hole 1 px | 0.501299376 | 0.504677755 |
| ser saddle 1 px,roundness | 0.504755717 | 0.504469854 |
| ser spot 1 px,ser hole 1 px | 0.504080042 | 0.504469854 |
| profile 1 2,ser spot 1 px | 0.504235967 | 0.504158004 |
| threshold compactness 60,radial mean | 0.508783784 | 0.504158004 |
| area m,width m | 0.503222453 | 0.504158004 |
| symmetry 03,symmetry 04 | 0.501559252 | 0.504054054 |
| ser spot 1 px,perimeter m | 0.506756757 | 0.503950104 |
| ser hole 1 px,perimeter m | 0.508316008 | 0.503950104 |
| symmetry 04,ser bright 1 px | 0.50524948 | 0.503950104 |
| symmetry 15,length m | 0.509225572 | 0.503846154 |
| symmetry 05,radial mean | 0.506730769 | 0.503846154 |
| symmetry 12,ser hole 1 px | 0.503586279 | 0.503742204 |
| ser ridge 1 px,width m | 0.505665281 | 0.503638254 |
| ser hole 1 px,ser bright 1 px | 0.505821206 | 0.503638254 |
| symmetry 12,ser bright 1 px | 0.505795218 | 0.503638254 |
| symmetry 02,ser hole 1 px | 0.503690229 | 0.503638254 |
| symmetry 14,ser bright 1 px | 0.504106029 | 0.503638254 |
| threshold compactness 60,ser bright 1 px | 0.505639293 | 0.503534304 |
| threshold compactness 50,radial mean | 0.508160083 | 0.503534304 |
| symmetry 05,ser bright 1 px | 0.503040541 | 0.503430353 |
| symmetry 13,ser hole 1 px | 0.50283264 | 0.503430353 |
| symmetry 04,ser hole 1 px | 0.503404366 | 0.503430353 |
| threshold compactness 60,perimeter m | 0.506704782 | 0.503326403 |
| symmetry 03,symmetry 05 | 0.503898129 | 0.503326403 |
| threshold compactness 40,radial mean | 0.50797817 | 0.503326403 |
| threshold compactness 40,ser bright 1 px | 0.505327443 | 0.503222453 |
| symmetry 02,ser bright 1 px | 0.506626819 | 0.503222453 |
| axial length ratio,perimeter m | 0.506133056 | 0.503118503 |
| threshold compactness 30,radial mean | 0.506860707 | 0.503118503 |
| symmetry 03,perimeter m | 0.507484407 | 0.503014553 |
| profile 2 2,perimeter m | 0.506470894 | 0.503014553 |
| symmetry 04,length m | 0.505847193 | 0.502910603 |
| symmetry 02,perimeter m | 0.505821206 | 0.502910603 |
| threshold compactness 50,perimeter m | 0.506340956 | 0.502806653 |
| threshold compactness 50,ser bright 1 px | 0.504989605 | 0.502806653 |
| radial mean,ser bright 1 px | 0.505483368 | 0.502598753 |
| symmetry 04,radial mean | 0.506340956 | 0.502494802 |
| symmetry 12,perimeter m | 0.505509356 | 0.502494802 |
| threshold compactness 40,perimeter m | 0.506418919 | 0.502390852 |
| threshold compactness 30,ser bright 1 px | 0.504963617 | 0.502390852 |
| roundness,perimeter m | 0.505613306 | 0.502286902 |

| | | |
|---|---|---|
| symmetry 15,radial mean | 0.506782744 | 0.502286902 |
| threshold compactness 30,perimeter m | 0.505899168 | 0.502286902 |
| symmetry 12,length m | 0.505535343 | 0.502182952 |
| symmetry 13,ser bright 1 px | 0.50493763 | 0.502182952 |
| profile 2 2,ser spot 1 px | 0.500233888 | 0.502182952 |
| symmetry 15,perimeter m | 0.506211019 | 0.502079002 |
| symmetry 05,axial length ratio | 0.502676715 | 0.502079002 |
| threshold compactness 40,ratio width to length | 0.504028067 | 0.502079002 |
| symmetry 03,radial mean | 0.507172557 | 0.501975052 |
| threshold compactness 30,width m | 0.502260915 | 0.501975052 |
| perimeter m,width m | 0.501975052 | 0.501975052 |
| symmetry 05,symmetry 12 | 0.501299376 | 0.501975052 |
| radial relative deviation,ser bright 1 px | 0.504132017 | 0.501871102 |
| axial small length,axial length ratio | 0.50472973 | 0.501871102 |
| ser bright 1 px,width m | 0.503456341 | 0.501767152 |
| threshold compactness 60,axial small length | 0.501689189 | 0.501663202 |
| symmetry 13,radial mean | 0.506574844 | 0.501559252 |
| symmetry 02,symmetry 04 | 0.499740125 | 0.501559252 |
| profile 2 2,width m | 0.49968815 | 0.501455301 |
| radial relative deviation,width m | 0.501065489 | 0.501351351 |
| threshold compactness 60,profile 2 2 | 0.500025988 | 0.501351351 |
| profile 1 2,perimeter m | 0.505301455 | 0.501247401 |
| symmetry 14,length m | 0.504209979 | 0.501247401 |
| threshold compactness 50,profile 2 2 | 0.5002079 | 0.501143451 |
| symmetry 03,symmetry 15 | 0.502234927 | 0.501143451 |
| threshold compactness 30,profile 2 2 | 0.500831601 | 0.501143451 |
| profile 1 2,width m | 0.500779626 | 0.501143451 |
| axial length ratio,radial mean | 0.506392931 | 0.501039501 |
| symmetry 15,profile 2 2 | 0.498570686 | 0.501039501 |
| threshold compactness 50,ser spot 1 px | 0.500233888 | 0.501039501 |
| symmetry 14,ser hole 1 px | 0.501481289 | 0.501039501 |
| symmetry 04,axial length ratio | 0.501689189 | 0.501039501 |
| symmetry 13,profile 2 2 | 0.497869023 | 0.501039501 |
| perimeter m,length m | 0.499428274 | 0.501039501 |
| symmetry 05,symmetry 13 | 0.49989605 | 0.500935551 |
| symmetry 14,axial length ratio | 0.501793139 | 0.500935551 |
| threshold compactness 40,profile 2 2 | 0.500493763 | 0.500831601 |
| axial length ratio,profile 2 2 | 0.497713098 | 0.500831601 |
| profile 1 2,ser bright 1 px | 0.504183992 | 0.500831601 |
| symmetry 12,axial length ratio | 0.501637214 | 0.500831601 |
| ser bright 1 px,perimeter m | 0.503742204 | 0.500831601 |
| symmetry 13,perimeter m | 0.506133056 | 0.500727651 |
| roundness,width m | 0.501845114 | 0.500727651 |
| symmetry 15,ser bright 1 px | 0.504703742 | 0.500727651 |
| threshold compactness 40,width m | 0.500259875 | 0.500727651 |
| symmetry 03,profile 2 2 | 0.496933472 | 0.500623701 |
| threshold compactness 30,ser saddle 1 px | 0.500545738 | 0.500623701 |

| | | |
|---|---|---|
| symmetry 02,axial length ratio | 0.501819127 | 0.500623701 |
| symmetry 03,symmetry 14 | 0.50031185 | 0.500623701 |
| axial length ratio,ser saddle 1 px | 0.500363825 | 0.500623701 |
| symmetry 13,symmetry 15 | 0.501741164 | 0.500519751 |
| symmetry 05,profile 2 2 | 0.498622661 | 0.500519751 |
| threshold compactness 40,ser saddle 1 px | 0.500389813 | 0.500519751 |
| axial small length,ser saddle 1 px | 0.500337838 | 0.5004158 |
| threshold compactness 60,area m | 0.5002079 | 0.5004158 |
| symmetry 04,perimeter m | 0.503976091 | 0.5004158 |
| symmetry 02,symmetry 13 | 0.500259875 | 0.5004158 |
| symmetry 12,profile 2 2 | 0.498570686 | 0.5004158 |
| ser valley 1 px,ratio width to length | 0.499376299 | 0.5004158 |
| axial small length,radial mean | 0.505613306 | 0.50031185 |
| symmetry 04,symmetry 12 | 0.499116424 | 0.50031185 |
| radial relative deviation,profile 2 2 | 0.498700624 | 0.50031185 |
| symmetry 13,symmetry 14 | 0.500675676 | 0.50031185 |
| profile 1 2,area m | 0.5002079 | 0.5002079 |
| threshold compactness 60,ser spot 1 px | 0.499922037 | 0.5002079 |
| symmetry 12,symmetry 13 | 0.499844075 | 0.5002079 |
| symmetry 03,axial length ratio | 0.501507277 | 0.5002079 |
| ser spot 1 px,area m | 0.50010395 | 0.50010395 |
| symmetry 15,ser saddle 1 px | 0.499922037 | 0.50010395 |
| radial relative deviation,area m | 0.500051975 | 0.50010395 |
| radial relative deviation,perimeter m | 0.504859667 | 0.50010395 |
| ser valley 1 px,area m | 0.500051975 | 0.50010395 |
| symmetry 12,area m | 0.50010395 | 0.50010395 |
| symmetry 04,profile 2 2 | 0.497843035 | 0.50010395 |
| threshold compactness 50,ser saddle 1 px | 0.500077963 | 0.50010395 |
| threshold compactness 40,ser spot 1 px | 0.499948025 | 0.50010395 |
| symmetry 04,ser saddle 1 px | 0.5002079 | 0.50010395 |
| symmetry 02,area m | 0.50010395 | 0.50010395 |
| radial mean,profile 2 2 | 0.495816008 | 0.50010395 |
| symmetry 03,area m | 0.50010395 | 0.50010395 |
| ser dark 1 px,area m | 0.500051975 | 0.50010395 |
| axial length ratio,area m | 0.500051975 | 0.50010395 |
| symmetry 05,area m | 0.50010395 | 0.50010395 |
| ser edge 1 px,area m | 0.5 | 0.50010395 |
| ser hole 1 px,area m | 0.50010395 | 0.50010395 |
| threshold compactness 50,axial small length | 0.501039501 | 0.5 |
| radial mean,width m | 0.501195426 | 0.5 |
| symmetry 14,ser saddle 1 px | 0.5 | 0.5 |
| symmetry 02,symmetry 14 | 0.499870062 | 0.5 |
| radial mean,length m | 0.500077963 | 0.5 |
| ser saddle 1 px,area m | 0.499844075 | 0.5 |
| ser bright 1 px,length m | 0.503976091 | 0.5 |
| profile 1 2,profile 2 2 | 0.498882536 | 0.49989605 |
| threshold compactness 40,area m | 0.499948025 | 0.49989605 |

| | | |
|---|---|---|
| threshold compactness 50,area m | 0.499948025 | 0.49989605 |
| symmetry 14,area m | 0.49989605 | 0.49989605 |
| profile 2 2,area m | 0.499948025 | 0.49989605 |
| ser bright 1 px,area m | 0.49989605 | 0.49989605 |
| symmetry 13,area m | 0.499844075 | 0.49989605 |
| symmetry 15,area m | 0.49989605 | 0.49989605 |
| ser ridge 1 px,area m | 0.49989605 | 0.49989605 |
| symmetry 12,symmetry 15 | 0.500051975 | 0.49989605 |
| threshold compactness 30,area m | 0.499948025 | 0.49989605 |
| symmetry 02,symmetry 15 | 0.499870062 | 0.49989605 |
| symmetry 05,ser saddle 1 px | 0.5002079 | 0.4997921 |
| symmetry 03,symmetry 12 | 0.499480249 | 0.4997921 |
| symmetry 04,area m | 0.49989605 | 0.4997921 |
| profile 2 2,ser edge 1 px | 0.502650728 | 0.4997921 |
| radial mean,area m | 0.4997921 | 0.4997921 |
| symmetry 02,ser saddle 1 px | 0.499870062 | 0.4997921 |
| ser dark 1 px,ratio width to length | 0.498934511 | 0.4997921 |
| axial small length,area m | 0.4997921 | 0.4997921 |
| radial relative deviation,ser saddle 1 px | 0.499818087 | 0.4997921 |
| symmetry 12,ser saddle 1 px | 0.499662162 | 0.49968815 |
| symmetry 04,width m | 0.500597713 | 0.49968815 |
| symmetry 14,perimeter m | 0.503222453 | 0.49968815 |
| symmetry 14,symmetry 15 | 0.499948025 | 0.49968815 |
| symmetry 03,ser saddle 1 px | 0.4997921 | 0.49968815 |
| symmetry 14,width m | 0.501377339 | 0.49968815 |
| symmetry 04,symmetry 13 | 0.499870062 | 0.49968815 |
| symmetry 13,ser saddle 1 px | 0.49989605 | 0.49968815 |
| threshold compactness 60,ser saddle 1 px | 0.499558212 | 0.4995842 |
| symmetry 03,width m | 0.499532225 | 0.4995842 |
| threshold compactness 30,ser spot 1 px | 0.499662162 | 0.4995842 |
| symmetry 02,profile 2 2 | 0.498804574 | 0.4995842 |
| profile 1 2,ser saddle 1 px | 0.499636175 | 0.4995842 |
| profile 2 2,ser saddle 1 px | 0.4997921 | 0.499480249 |
| symmetry 14,profile 2 2 | 0.497401247 | 0.499480249 |
| symmetry 04,symmetry 15 | 0.499532225 | 0.499480249 |
| symmetry 15,width m | 0.500935551 | 0.499376299 |
| radial mean,ser saddle 1 px | 0.499298337 | 0.499376299 |
| profile 1 2,ser hole 1 px | 0.498388773 | 0.499272349 |
| threshold compactness 30,ratio width to length | 0.499948025 | 0.499272349 |
| symmetry 02,width m | 0.499948025 | 0.499168399 |
| symmetry 12,width m | 0.500363825 | 0.499168399 |
| symmetry 05,symmetry 15 | 0.500519751 | 0.499064449 |
| symmetry 04,symmetry 05 | 0.501923077 | 0.499064449 |
| symmetry 03,ser edge 1 px | 0.499324324 | 0.499064449 |
| ser ridge 1 px,perimeter m | 0.502130977 | 0.499064449 |
| ser ridge 1 px,ser saddle 1 px | 0.499740125 | 0.498960499 |
| profile 1 2,ser edge 1 px | 0.498570686 | 0.498648649 |

| | | |
|---|---|---|
| ser ridge 1 px,ser valley 1 px | 0.498544699 | 0.498544699 |
| ser ridge 1 px,mito aspect ratio ar | 0.499194387 | 0.498440748 |
| threshold compactness 40,axial small length | 0.498856549 | 0.498336798 |
| axial small length,perimeter m | 0.503118503 | 0.498232848 |
| threshold compactness 50,width m | 0.498778586 | 0.498232848 |
| symmetry 05,symmetry 14 | 0.498986486 | 0.498232848 |
| symmetry 02,length m | 0.501507277 | 0.498232848 |
| threshold compactness 30,axial length ratio | 0.499766112 | 0.498128898 |
| threshold compactness 30,axial small length | 0.497141372 | 0.497817048 |
| symmetry 05,ser edge 1 px | 0.498154886 | 0.497817048 |
| ser edge 1 px,ratio width to length | 0.497089397 | 0.497713098 |
| threshold compactness 40,axial length ratio | 0.500129938 | 0.497713098 |
| threshold compactness 60,width m | 0.497765073 | 0.497609148 |
| ser ridge 1 px,ser dark 1 px | 0.496855509 | 0.497401247 |
| axial small length,ser hole 1 px | 0.497765073 | 0.497401247 |
| symmetry 14,axial small length | 0.499012474 | 0.497297297 |
| axial length ratio,width m | 0.497531185 | 0.497297297 |
| symmetry 03,symmetry 13 | 0.498986486 | 0.497297297 |
| radial relative deviation,ser edge 1 px | 0.496985447 | 0.497193347 |
| axial small length,profile 2 2 | 0.49454262 | 0.497193347 |
| symmetry 14,ser edge 1 px | 0.496881497 | 0.496881497 |
| ser spot 1 px,width m | 0.498960499 | 0.496777547 |
| axial small length,ser valley 1 px | 0.496699584 | 0.496465696 |
| threshold compactness 50,axial length ratio | 0.499428274 | 0.496465696 |
| profile 1 2,length m | 0.494386694 | 0.496257796 |
| ser ridge 1 px,length m | 0.49716736 | 0.496257796 |
| symmetry 12,axial small length | 0.498960499 | 0.496153846 |
| symmetry 13,width m | 0.499844075 | 0.496153846 |
| symmetry 02,axial small length | 0.498648649 | 0.496049896 |
| symmetry 03,axial small length | 0.496595634 | 0.495841996 |
| symmetry 04,axial small length | 0.497323285 | 0.495841996 |
| ser spot 1 px,ser saddle 1 px | 0.49516632 | 0.495426195 |
| threshold compactness 60,axial length ratio | 0.498830561 | 0.495218295 |
| symmetry 12,threshold compactness 60 | 0.497037422 | 0.494386694 |
| symmetry 12,ser edge 1 px | 0.49254158 | 0.494282744 |
| symmetry 02,ser edge 1 px | 0.493970894 | 0.494282744 |
| symmetry 15,ser edge 1 px | 0.495114345 | 0.493970894 |
| symmetry 04,threshold compactness 60 | 0.496387734 | 0.493451143 |
| radial relative deviation,length m | 0.498128898 | 0.493451143 |
| symmetry 14,threshold compactness 60 | 0.496517672 | 0.493451143 |
| symmetry 05,width m | 0.50010395 | 0.493243243 |
| axial length ratio,ser edge 1 px | 0.494776507 | 0.492411642 |
| symmetry 15,ser valley 1 px | 0.493165281 | 0.491995842 |
| ser bright 1 px,ser dark 1 px | 0.491034304 | 0.491580042 |
| symmetry 15,axial small length | 0.494672557 | 0.491372141 |
| ser saddle 1 px,ratio width to length | 0.491268191 | 0.491164241 |
| symmetry 13,ser edge 1 px | 0.492983368 | 0.490956341 |

| | | |
|---|---|---|
| symmetry 15,threshold compactness 60 | 0.491865904 | 0.490748441 |
| radial mean,ser hole 1 px | 0.489760915 | 0.48970894 |
| radial relative deviation,ser valley 1 px | 0.488539501 | 0.489397089 |
| symmetry 04,symmetry 14 | 0.488071726 | 0.488357588 |
| symmetry 02,threshold compactness 60 | 0.49002079 | 0.487422037 |
| symmetry 15,ser dark 1 px | 0.486382536 | 0.487006237 |
| threshold compactness 60,ser edge 1 px | 0.485758836 | 0.487006237 |
| symmetry 05,axial small length | 0.487889813 | 0.485446985 |
| symmetry 14,ser dark 1 px | 0.484953222 | 0.484615385 |
| symmetry 05,ser dark 1 px | 0.48497921 | 0.484511435 |
| profile 1 2,ser valley 1 px | 0.485317048 | 0.484511435 |
| threshold compactness 30,ser edge 1 px | 0.483549896 | 0.484303534 |
| symmetry 14,ser valley 1 px | 0.485914761 | 0.483367983 |
| symmetry 05,threshold compactness 60 | 0.482926195 | 0.482848233 |
| symmetry 03,threshold compactness 60 | 0.482744283 | 0.482848233 |
| symmetry 13,threshold compactness 60 | 0.482926195 | 0.482848233 |
| symmetry 04,ser dark 1 px | 0.480067568 | 0.482432432 |
| area m,perimeter m | 0.488565489 | 0.481496881 |
| threshold compactness 40,ser edge 1 px | 0.481133056 | 0.481496881 |
| symmetry 04,ser valley 1 px | 0.481107069 | 0.47962578 |
| symmetry 04,ser edge 1 px | 0.481211019 | 0.47952183 |
| threshold compactness 50,ser edge 1 px | 0.480509356 | 0.479417879 |
| threshold compactness 30,ser dark 1 px | 0.476793139 | 0.477027027 |
| threshold compactness 40,ser dark 1 px | 0.476247401 | 0.476299376 |
| threshold compactness 50,ser dark 1 px | 0.476559252 | 0.476195426 |
| threshold compactness 30,ser valley 1 px | 0.474194387 | 0.474636175 |
| threshold compactness 40,ser valley 1 px | 0.474090437 | 0.474532225 |
| ser spot 1 px,ser edge 1 px | 0.475441788 | 0.474116424 |
| symmetry 13,ser valley 1 px | 0.472349272 | 0.473908524 |
| threshold compactness 60,ser dark 1 px | 0.473804574 | 0.473804574 |
| symmetry 12,ser valley 1 px | 0.473362786 | 0.473284823 |
| symmetry 13,axial small length | 0.473336798 | 0.473180873 |
| symmetry 13,ser dark 1 px | 0.472453222 | 0.473076923 |
| radial relative deviation,ser dark 1 px | 0.474350312 | 0.472557173 |
| threshold compactness 50,ser valley 1 px | 0.472609148 | 0.472245322 |
| threshold compactness 60,ser valley 1 px | 0.472323285 | 0.471933472 |
| axial small length,ser dark 1 px | 0.472713098 | 0.470997921 |
| symmetry 12,ser dark 1 px | 0.469360707 | 0.46954262 |
| ser spot 1 px,ser dark 1 px | 0.46964657 | 0.469022869 |
| axial length ratio,ser dark 1 px | 0.469048857 | 0.469022869 |
| symmetry 02,ser dark 1 px | 0.469100832 | 0.468814969 |
| ser spot 1 px,ser valley 1 px | 0.468269231 | 0.467775468 |
| profile 1 2,ser dark 1 px | 0.467151767 | 0.467255717 |
| symmetry 02,ser valley 1 px | 0.467073805 | 0.466943867 |
| symmetry 03,ser dark 1 px | 0.466268191 | 0.466839917 |
| profile 2 2,ser dark 1 px | 0.463721414 | 0.464345114 |
| axial length ratio,ser valley 1 px | 0.463201663 | 0.463617464 |

| | | |
|---|---|---|
| profile 2 2,ser valley 1 px | 0.462967775 | 0.462785863 |
| symmetry 03,ser valley 1 px | 0.462032225 | 0.462785863 |
